# Battle for the histones: a secreted bacterial sirtuin from *Campylobacter jejuni* activates neutrophils and induces inflammation during infection

**DOI:** 10.1101/2022.07.19.497369

**Authors:** Sean M. Callahan, Trevor J. Hancock, Ryan S. Doster, Caroline B. Parker, Mary E. Wakim, Jennifer A. Gaddy, Jeremiah G. Johnson

**Affiliations:** Department of Microbiology, University of Tennessee, Knoxville, TN 37996, USA; Division of Infectious Diseases, Department of Medicine Vanderbilt University Medical Center, Nashville, TN 37232, USA

**Author notes:** **Corresponding Author:** Jeremiah G. Johnson, MS, PhD, 516 Ken and Blaire Mossman Bldg., 1311 Cumberland Avenue, Knoxville, TN 37996.

**Keywords:** *Campylobacter jejuni*, neutrophils, histones, sirtuins, deacetylation, inflammation, HDAC, host-microbe interaction

## Abstract

Histone modifications alter numerous cornerstone processes in eukaryotes, including metabolism, physiology, and immunity. Numerous bacterial pathogens can alter expression of host-derived sirtuins to deacetylate histones in order to promote infection, yet, a bacterial-derived sirtuin has yet to be investigated to deacetylate host histones. Using *Campylobacter jejuni*, the leading cause of bacterial-derived gastroenteritis, we found a secreted sirtuin, SliP, which binds to and deacetylates neutrophil histones. We found neutrophil activation and extrusion of neutrophil extracellular traps was SliP dependent, whereby *sliP* mutants are unable to activate neutrophils or promote NETosis. Leveraging the mouse model of campylobacteriosis, we further demonstrate the *sliP* mutant can efficiently infect IL-10^-/-^ mice, but induction of proinflammatory cytokine production and gastrointestinal pathology is SliP-dependent. In conclusion, we investigate a unique bacterial effector which targets host histones and is responsible for the inflammatory response and tissue pathology observed during campylobacteriosis.

**Highlights:**

- *C. jejuni* encodes a secreted effector, SliP, which functions as a canonical sirtuin
- SliP binds to and deacetylates neutrophil histone H3 during bacterial infection
- *C. jejuni*-induced neutrophil activation and NETosis are SliP-dependent
- Inflammation and tissue pathology during *C. jejuni* infection is SliP-dependent

## Main

*Campylobacter* species are the leading cause of bacterial-derived gastroenteritis worldwide, infecting a projected ninety million individuals (Man, 2011; World Health Organization, 2015). While infection is often self-limiting in immunocompetent individuals, numerous post-infectious disorders can develop, including Guillain-Barré syndrome (GBS), irritable bowel syndrome (IBS), and reactive arthritis (ReA) (Callahan et al., 2021a; Reti et al., 2015). In addition to the prevalence of infection and these various outcomes, the high incidence of antibiotic resistance has led both the Centers for Disease Control and World Health Organization to classify *Campylobacter* spp. as serious threats to public health (Silva et al., 2011; Young et al., 2007). Despite these impacts, little is known about what bacterial and host factors are responsible for the inflammation during human campylobacteriosis and how that might be responsible for intestinal disease. In our previous work, we found neutrophils are recruited to the site of infection and that predominantly neutrophil-derived proteins are present in the fecal samples of *C. jejuni*-infected patients. In addition, we found *C. jejuni* is a potent activator of human neutrophils and stimulation leads to the production of neutrophil extracellular traps (NETs), which appear to localize with abscesses in colonic crypts (Callahan et al., 2020; Shank et al., 2018). Neutrophil activation within campylobacteriosis has been associated with pro-inflammatory and tissue pathology (Sørensen et al., 2013; Sun et al., 2012, 2013), therefore, understanding the microbial factors that lead to this could lead to the development of novel therapeutics. To identify bacterial factors which promote neutrophil activation by *C. jejuni*, including the production of NETs, we screened a transposon library of *C. jejuni* for mutants unable to activate primary human neutrophils and identified a locus (Cjj81176_0779) that encodes a putative bacterial sirtuin.

Post-translational modifications (PTMs) in eukaryotes play a vital role in orchestrating host-microbe interactions. Differential acetylation of host proteins is accomplished using numerous enzymes, including histone acetyltransferases (HATs), histone deacetylases (HDACs), and sirtuins. The outcomes of altering host protein acetylation are varied and include impacts to processes involved in autoimmunity, inflammation, and tumorigenesis (Barneda-Zahonero and Parra, 2012; Hamminger et al., 2020; Shakespear et al., 2011). Because the host uses acetylation to control these cellular processes, pathogens, including *Listeria monocytogenes, Salmonella typhimurium*, and *Mycobacterium tuberculosis*, can manipulate host proteins to promote infection (Bhaskar et al., 2020; Gogoi et al., 2018; Pennini et al., 2010; Pereira et al., 2018; Ribet and Cossart, 2010a, 2010b; Rolando et al., 2013). However, recent research has found that bacterial-derived effectors can directly modify host proteins PTMs. While some pathogens have acquired and incorporated host enzymes into their virulence factor repertoire, others have convergently evolved to mimic the activity of host enzymes (Bernal et al., 2014; Luu and Carabetta, 2021; Salomon and Orth, 2013). Importantly, bacterial effectors that are translocated and directly bind host proteins to differentially acetylate them remain to be identified.

In eukaryotes, a common target of PTMs are histones. Histones play a critical role in the structure and function of chromatin, which have profound effects on gene expression. Core histones are characterized by a histone folding domain and N-terminal tails accessible to enzymes involved in PTMs (Mariño-Ramírez et al., 2005). For example, HATs acetylate the *ε*-amino group of lysine residues on histones, which results in a neutral charge and therefore opens the chromatin for RNA polymerase (Glozak and Seto, 2007). In contrast, during histone deacetylation, the acetyl group is removed from lysine residues, which causes the chromatin to close and prevent RNA polymerase binding (Kurdistani and Grunstein, 2003). Beyond reducing gene expression, other research found HDACs can positively regulate gene expression by limiting acetylation near gene bodies and intergenic regions, which instead allows RNA elongation factors to bind more readily (Greer et al., 2015; Haberland et al., 2009; Nusinzon and Horvath, 2005). As a result, numerous pro-inflammatory genes are upregulated in response to HDAC activity, including IL-8, CSF-1, and TNF-α (Gatla et al., 2017; Marumo et al., 2010; Zhu et al., 2010).

In addition to changing gene expression, differential acetylation of proteins, including histones, can lead to other cellular responses. In neutrophils, for example, increased HDAC activity promotes citrullination of histone H3 by protein arginine deiminase 4 (PAD4), which results in the elaboration of neutrophil extracellular traps (NETs) and the associated inflammation (Chen et al., 2020; Hamam and Palaniyar, 2019; Poli et al., 2021). This post-translational conversion of arginine to citrulline on histone H3 through PAD4 replaces the primary ketimide with a ketone group and is indispensable for the production of NETs (Rohrbach et al., 2012). Further, PAD4 promotes decondensation of heterochromatin to facilitate NET release into the extracellular milieu (Leshner et al., 2012). Despite the apparent relation between histone acetylation and neutrophil physiology, the role of sirtuins in these responses are not well defined.

Importantly, because histone acetylation plays a role in proinflammatory gene expression and cellular outcomes, investigators have sought to characterize their effects in various diseases and whether altering these responses can improve health. For example, in the gastrointestinal tract, deacetylation inhibition is a naturally occurring phenomenon. Sodium butyrate is a short-chain fatty acid (SCFA) produced by the human gut microbiota as a byproduct of anaerobic fermentation of dietary fibers (Silva et al., 2020). By binding to and inhibiting the enzymatic activity of various HDACs and sirtuins, butyrate has been shown to reduce inflammation within the gastrointestinal tract, specifically through suppressing neutrophil activity and inhibiting nuclear factor κB (NF-κB)-dependent proinflammatory gene expression in colonocytes (Canani, 2011; Siddiqui and Cresci, 2021; Yin et al., 2001). Similarly, trichostatin A (TSA) is a potent HDAC inhibitor and was recently discovered to also interact with mammalian sirtuins (Vigushin et al., 2001; You and Steegborn, 2018). Similarly to sodium butyrate, TSA suppresses cellular inflammatory responses and promotes neutrophil apoptosis (Kankaanranta et al., 2010; Toki et al., 2016). Because of these results, HDACs and sirtuins are emerging therapeutic targets for the development of inhibitors that can treat inflammatory disorders of the gastrointestinal tract.

In the present study, we characterize a novel bacterial effector protein in *C. jejuni*, which we have termed SliP, which we identified as being involved in the activation of human neutrophils and the production of NETs. We show SliP functions as an NAD^+^ and zinc-dependent sirtuin which deacetylates acetylated peptides and is secreted into neutrophils using the flagellar secretion apparatus. Following secretion, SliP associates with neutrophil histones, which correlates with a reduction in the acetylation of these structures. Finally, we demonstrate inflammation and intestinal pathology is promoted by SliP despite the mutant being able to effectively colonize and disseminate within the host. These findings suggest histone deacetylation mediated by SliP is a novel mechanism by which a pathogen induces an inflammatory response and tissue pathology during infection. With these findings, we expect therapeutic strategies can be developed to mitigate SliP-dependent inflammation and pathology during acute and chronic human disease.

## Results

### *C. jejuni* encodes a sirtuin-like deacetylase more frequently present in clinical isolates than agricultural or environmental isolates

The DUF4917 family of proteins are ubiquitous in microorganisms, with many representatives found in *Burkholderia* and *Brucella* species, but little is known about the function of these proteins (EMBL-EFI). The Cjj81176_0779 locus we identified in *C. jejuni* is predicted to contain a DUF4917 domain with shared amino acid homology with other human bacterial pathogen-derived DUF4917 family proteins, including those from *Yersinia enterocolitica* and *Legionella pneumophila* (Figure 1A) (Milne-Davies et al., 2019; Shames et al., 2017). As this family of proteins has yet to be investigated for functionality, secondary structure prediction was performed using Phyre2 and iTASSER (Kelley et al., 2015; Roy et al., 2010). In both analyses, the protein encoded by Cjj81176_0779 appears to share secondary structure homology with bacterial sirtuins, specifically sirtuin-2, allowing us to name this protein SliP for **<u>s</u>**irtuin-**<u>li</u>**ke **<u>p</u>**rotein (Figure 1B). Important to this analysis, sirtuins contain a Rossmann fold comprised of a Gly-X-Gly sequence responsible for the binding of the phosphate group of NAD^+^ and a small pocket of charged amino acids which bind the two ribose molecules of NAD^+^ (Satoh et al., 2011). SliP possesses a Gly-Asn-Gly sequence and when the amino acid composition of SliP was compared to those of other bacterial and human sirtuins, the glycine residue at position 26 (G26) was found to be evolutionarily conserved in all sirtuins, which encodes the second glycine residue (Figure 1C, D; Supplemental Movie 1). To determine the potential role of this protein in the pathogenesis of *C. jejuni*, we examined for *sliP* in the genomes of agricultural and environmental isolates of *C. jejuni* and compared incidence to human clinical isolates collected and sequenced in a previous study (Kelley et al., 2020). *sliP* is significantly enriched in clinical isolates, with 36 of 70 isolates encoding *sliP* (Figure 1E) and only 3 of 66 agricultural and environmental *C. jejuni* possessing the gene (χ = 36.51, z = 6.042, p = < 0.0001).

**Figure 1.**
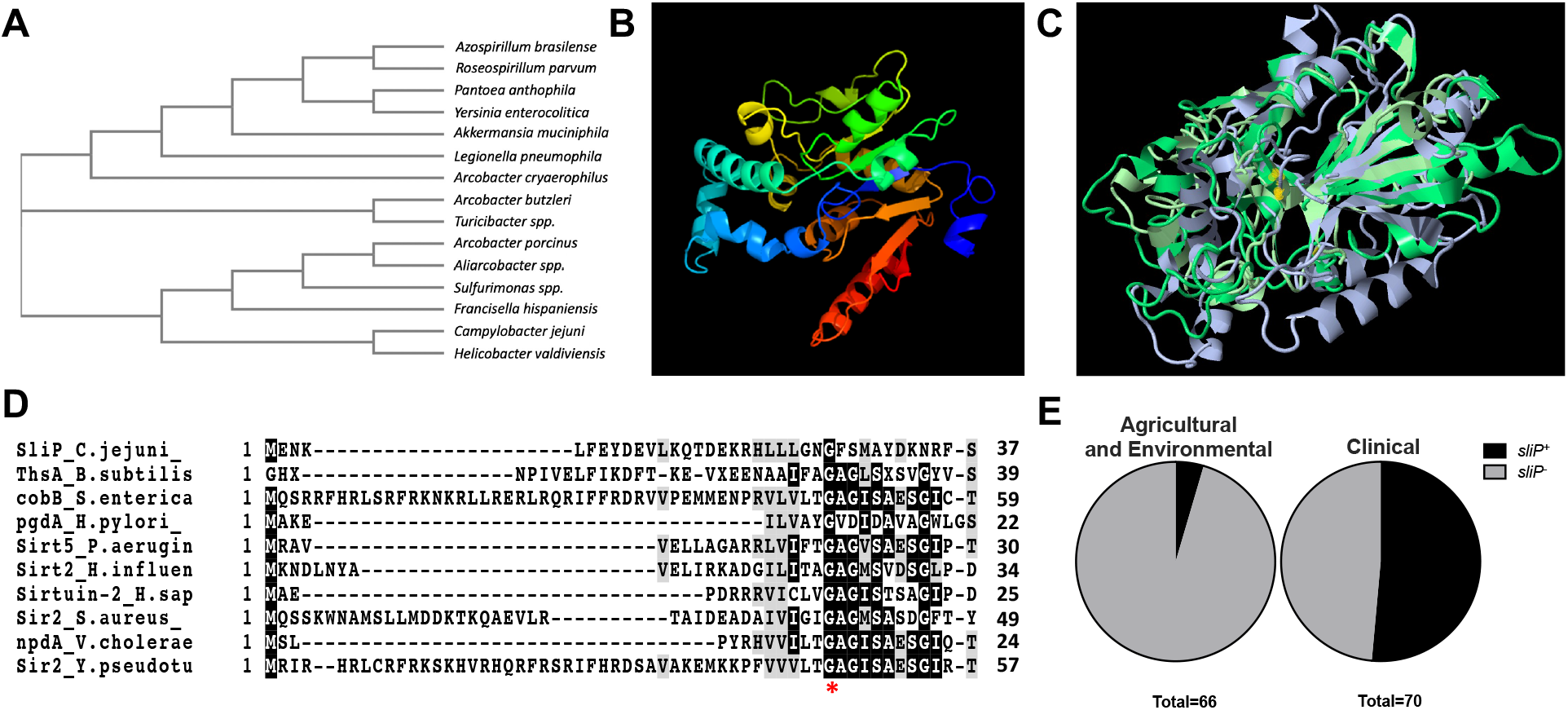
Identification of a *C. jejuni* sirtuin, SliP. (A) Phylogenetic analysis of the DUF4917 family proteins in relation to *C. jejuni*. A BLAST analysis was performed for proteins homologous to the *C. jejuni* DUF4917 family protein. (B) Prediction of the secondary structure of SliP using Phyre2. ((C) Overlapped secondary structure models of DUF4917 family proteins using Jmol software. Yellow dots indicate position of the G26 residue within the structural model for SliP. (D) Multiple sequence alignment of bacterial and human sirtuins, along with *C. jejuni* SliP. Amino acid sequences were pulled from NCBI and aligned using T-Coffee. The red star indicates the sole amino acid conserved in all sirtuins. (E) Percentage of *sliP* containing environmental and clinical *C. jejuni* strains from a previous study. Whole genomes were uploaded to KBase, where the homology threshold was set to 75%. Chi-square contingency testing was performed, χ^2^ = 36.51, z = 6.042, p = <0.0001.

### SliP functions as a canonical sirtuin by deacetylating lysines in an NAD^+^ and zinc-dependent manner

To investigate the function of SliP, His-tagged protein was purified by Ni-NTA chromatography (Figure S1A, B). Because NAD^+^ and zinc are often required for sirtuin activity (Moniot et al., 2012), the deacetylase activity of purified SliP was determined for acetyl-lysine peptides with or without those cofactors. As expected, when either 5 mM NAD^+^ or 20 μM zinc chloride were individually added to deacetylase reactions, there was a 1.92 and 3.11-fold increase in lysine deacetylation after one hour relative to reactions without the cofactors, respectively. When both NAD^+^ and zinc chloride were added, we observed a 3.76-fold increase in lysine deacetylation when compared to reactions without the cofactors (Figure 2A). As NAD^+^ dependent sirtuins classically hydrolyze NAD^+^ during lysine deacetylation (Imai and Guarente, 2014), we sought to determine whether SliP-mediated deacetylation leads to NAD^+^ hydrolysis. Using purified SliP, NAD^+^ concentrations were monitored over time during lysine deacetylation using a commercially available kit. As expected, NAD^+^ concentrations decreased 1.83-fold during deacetylation of acetyl-lysine peptides by purified SliP, while similarly purified lysates without recombinant SliP (empty vector) were unable to significantly decrease NAD^+^ concentrations (1.06-fold reduction) (Figure 2B). Importantly, deacetylase reactions containing SliP or empty vector lysates did not result in significant hydrolysis of NADH (Figure S2). To determine whether the conserved glycine residue in SliP (Figure 1C) is required for full sirtuin activity, a single point mutation was introduced into SliP (SliP_G26A_) (Figure S1C). Compared to wild-type SliP, SliP_G26A_ was unable to deacetylate lysine residues and hydrolyze NAD^+^ as readily, exhibiting 2.05-fold decrease and 1.29-fold increase, respectively (Figure 2B, C). As such, we hypothesize SliP functions as a canonical sirtuin, and the highly conserved glycine residue is involved in NAD^+^ binding and hydrolysis during lysine deacetylation.

**Figure 2.**
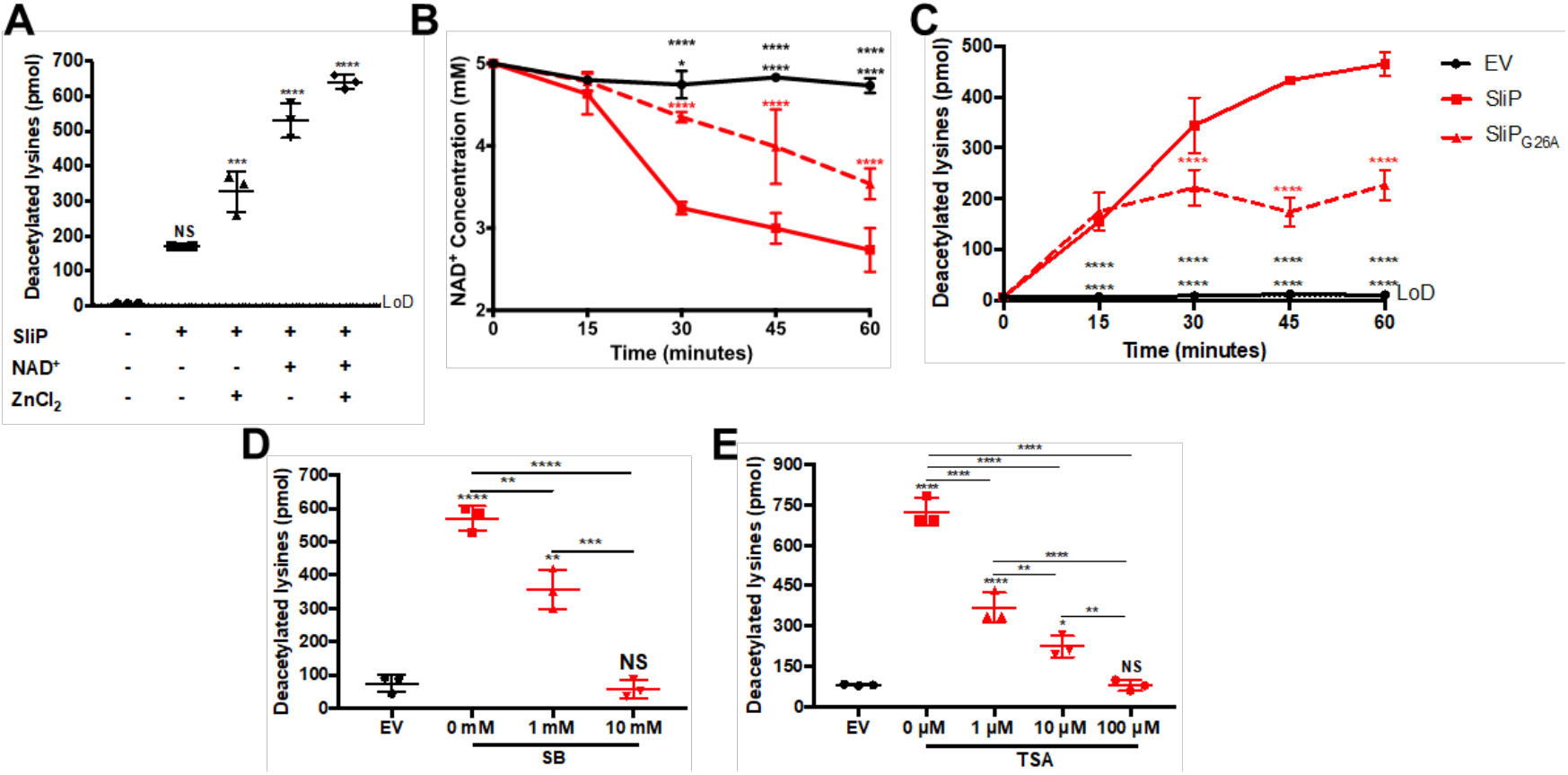
SliP is an NAD^+^ and zinc-dependent sirtuin that can be chemically inhibited by HDAC inhibitors. (A) Lysine deacetylation assay was performed for SliP with common sirtuin cofactors, NAD^+^ and zinc chloride one-hour post incubation at 37°C. For statistical analysis, each treatment group was compared back to the empty vector control. (B) NAD^+^ hydrolysis assay during (C) lysine deacetylation during a one-hour time course at 37°C. Black bars represent the empty vector (EV) control, whereby the SliP and SliP_G26A_ incubated reactions are represented in red solid and dashed lines, respectively. (D) Sodium butyrate (SB) and (E) trichostatin-A (TSA) dose-dependent chemical inhibition of SliP one-hour post incubation at 37°C. Multiple comparison testing was performed using ANOVA with Tukey’s *post-hoc* test. \**p* < .05; \*\**p* < .01; \*\*\**p* < .001; \*\*\*\**p* < .0001

As SliP appears to function as a sirtuin, we further sought to determine if characterized deacetylase inhibitors can reduce SliP activity. To accomplish this, we used conditions where maximum lysine deacetylation was observed and titrated in either media alone, sodium butyrate (SB), or trichostatin A (TSA) and assayed for lysine deacetylation as above. We observed a dose-dependent inhibition of deacetylation for both SB and TSA to where at 10 mM (for SB) and 100 μM (for TSA), the presence of SliP did not lead to significantly more deacetylation than when eluates lacking a deacetylase were used. For example, there was a 9.85-fold reduction in lysine deacetylation at 10 mM SB (Figure 2D) and a 9.01-fold reduction in lysine deacetylation at 100 μM TSA (Figure 2E). These results demonstrate SB and TSA can chemically inhibit SliP-dependent lysine deacetylation, which supports the conclusion that SliP functions as a canonical sirtuin.

### SliP is a secreted bacterial sirtuin which requires the flagellar secretion apparatus

To determine whether host proteins can be targeted by SliP during infection, we first established whether SliP was secreted from *C. jejuni*. To examine SliP secretion, polyclonal antibodies were raised against purified His-tagged *C. jejuni* SliP and cell-free supernatants were assayed for the presence of the protein by western blot. In addition, because deoxycholate has been shown to promote secretion by *C. jejuni* through the flagellar secretion system (Christensen et al., 2009; Konkel et al., 2004), we investigated whether SliP translocation into culture supernatants increased in response to sodium deoxycholate and whether SliP secretion was absent in a non-flagellar mutant of *C. jejuni* (Δ*flgE*). Regardless of whether strains were grown with or without sodium deoxycholate, SliP was detected in whole cell lysates of each strain apart from Δ*sliP* (Figure 3A). When we examined for translocation of SliP into the media lacking sodium deoxycholate, we did not detect SliP in those cell-free supernatants. In contrast, when strains were grown in media containing sodium deoxycholate, SliP was detected in the cell-free supernatants of all strains except the Δ*sliP* and Δ*flgE* mutants (Figure 3A). To eliminate the possibility that SliP was present in the supernatant due to sodium deoxycholate-induced cell lysis, we quantified the amount of *C. jejuni* genomic DNA in supernatants with or without sodium deoxycholate by qPCR and observed no significant differences across strains or growth conditions (Figure S3). This result supports our earlier observation that SliP is translocated through the flagellar secretion apparatus in response to an intestinal signal, such as deoxycholate.

**Figure 3.**
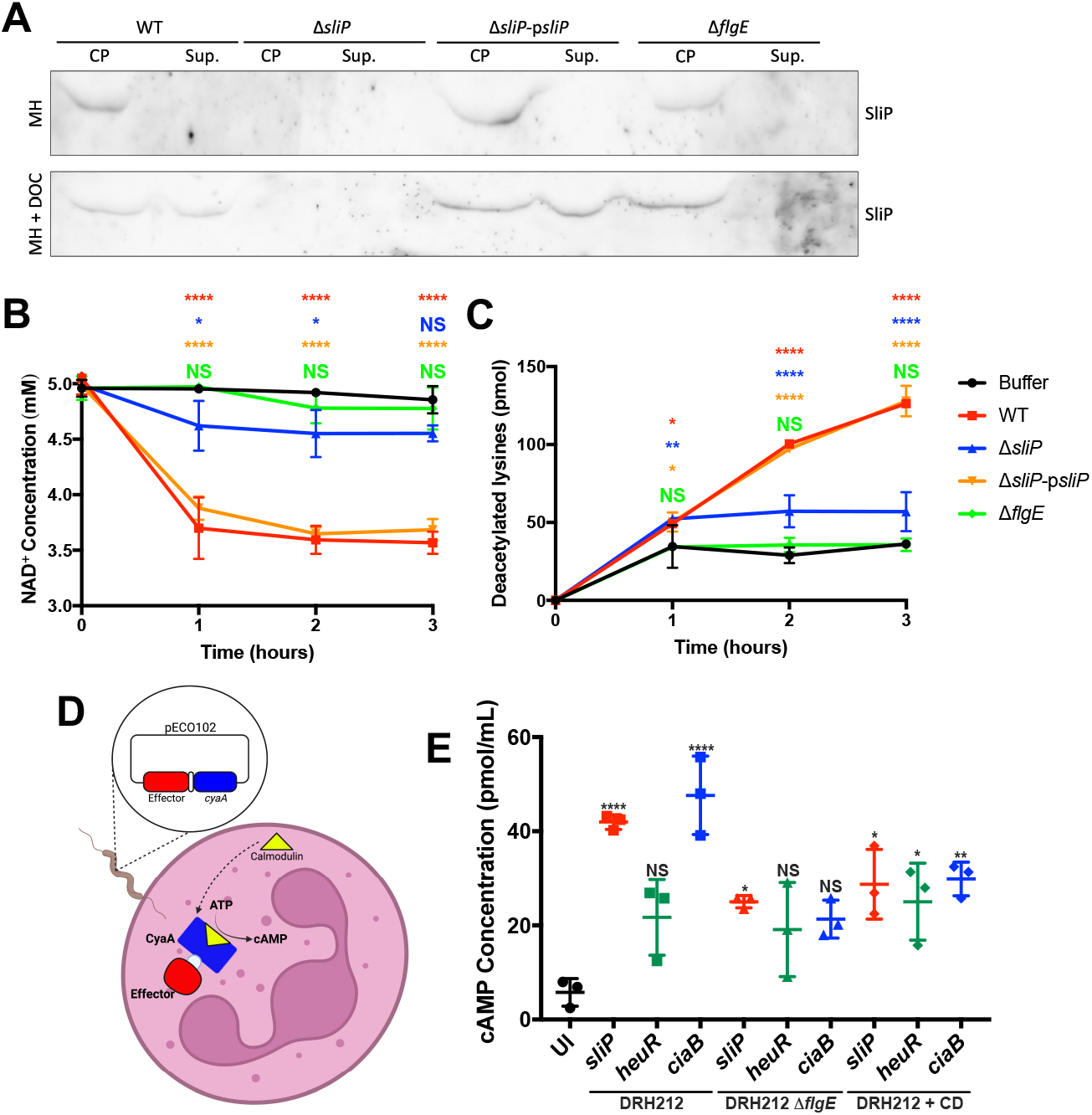
SliP is a secreted sirtuin that is translocated into host neutrophils through the flagellar secretion system. (A) Deoxycholate (DOC) induces *C. jejuni* SliP flagellar secretion into the supernatant. Cultures were grown for 48 hours under microaerobic conditions, whereby after incubation, supernatant was protein precipitated using TSA. Whole cell lysates and supernatants were analyzed for SliP abundance through western blot analysis. (B) *C. jejuni* supernatant is able to perform NAD^+^ hydrolysis during (C) Lysine deacetylation during a one-hour incubation period at 37°C. (D) Schematic for the calmodulin-dependent *B. pertussis* adenylate cyclase (CyaA) bacterial secretion assay within neutrophils. (E) CyaA translocation assay performed in wild-type and the flagella mutant (Δ*flgE*) infected neutrophils. The *ciaB-cyaA* and *heuR-cyaA* translational fusions served as positive and negative controls, respectively. Purified neutrophils were incubated with *C. jejuni* strains at an MOI of 10 for three hours before measurement of intracellular cAMP. Multiple comparison testing was performed using ANOVA with Tukey’s *post-hoc* test. \**p* < .05; \*\**p* < .01; \*\*\**p* < .001; \*\*\*\**p* < .0001

To determine whether secreted SliP is functional, cell-free supernatants of cultures grown in media supplemented with sodium deoxycholate were investigated for sirtuin activities, including lysine deacetylation and NAD^+^ hydrolysis. We observed significant NAD^+^ hydrolysis in supernatants from both wild-type and *sliP* complemented strains, with 26.52% and 24.12% reductions in NAD^+^ concentrations relative to the media alone control, respectively. In contrast, supernatants from both Δ*sliP* and flagellar hook mutant (Δ*flgE*) mutants did not produce significant NAD^+^ hydrolysis with only 6.24% and 1.61% reductions relative to the media alone control, respectively. Furthermore, we observed deacetylase activity within the supernatants of wild-type and *sliP* complemented strains, with the amount of lysines deacetylated increasing by 3.49 and 3.54-fold when compared to the media control, respectively. As expected, supernatants from the Δ*sliP* and Δ*flgE* mutants were unable to significantly deacetylate lysine residues, exhibiting 1.58 and 0.99-fold increases when compared to the media control, respectively (Figures 3B, C). This observation not only confirms the presence of SliP in *C. jejuni* supernatants, but also is enzymatically active as a sirtuin and requires the flagellar secretion apparatus for translocation. As such, we hypothesized SliP is a secreted bacterial effector which deacetylates lysine residues.

### SliP is secreted into human neutrophils and promotes elevated deacetylase activity

To investigate whether SliP is secreted into human neutrophils, we constructed plasmids containing translational fusions of proteins of interest to the *Bordetella pertussis* adenylate cyclase (CyaA) domain (Chakravarthy et al., 2017) (Figure 3D). In addition to the SliP-CyaA reporter, we also constructed HeuR-CyaA and CiaB-CyaA reporters as negative and positive secretion controls, respectively (Christensen et al., 2009; Johnson et al., 2016; Konkel et al., 1999). These plasmids were introduced into wild-type and Δ*flgE* strains as the flagellar T3SS is predicted to serve as the major secretion system in *C. jejuni* (Aizawa, 2001; Neal-McKinney and Konkel, 2012). Three hours post-infection, cAMP levels were measured as a proxy for effector secretion into host cells using an enzyme-linked immunosorbent assay (ELISA). In wild-type strains, cAMP levels increased by 7.26, 3.76, and 8.24-fold when they possessed the SliP-CyaA, HeuR-CyaA, and CiaB-CyaA plasmid, respectively, when compared to uninfected neutrophils. In contrast, when Δ*flgE* containing the same constructs were used to infect neutrophils, cAMP levels decreased by 1.68, 1.13, and 2.23-fold when compared to their wild-type SliP-CyaA, HeuR-CyaA, and CiaB-CyaA counterpart, respectively. Furthermore, strains encoding the various CyaA reporters were able to internalize at similar rates and produced non-significantly different intracellular CyaA despite the fusion (Figure S4A, B). Because decreased translocation of bacterial-derived proteins in Δ*flgE* could be due to reduced outside-to-inside secretion or endocytosis, actin rearrangement was blocked using cytochalasin D (CD). Treatment with CD resulted in reduced cAMP by 1.46, 0.87, and 1.6-fold when compared to untreated infections with wild-type SliP-CyaA, HeuR-CyaA, and CiaB-CyaA strains, respectively (Figure 3E).

Based on this data, we next sought to establish whether translocation of SliP is associated with intracellular environments which exhibit elevated lysine deacetylase activity. To accomplish this, human neutrophils were infected with wild-type *C. jejuni, ΔsliP, ΔsliP* complemented with the wild-type *sliP* allele (Δ*sliP*-p*sliP*), and Δ*sliP* complemented with the G26A substitution (Δ*sliP*-p*sliP*_G26A_). As predicted, intracellular lysine deacetylation was elevated in neutrophils infected with wild-type *C. jejuni* and Δ*sliP*-p*sliP;* however, neutrophils infected with Δ*sliP* possessed intracellular environments that exhibited significantly reduced lysine deacetylase activity despite increased uptake of the Δ*sliP* strain into neutrophils (Figure 4A; Figure S5). Furthermore, Δ*sliP* expressing the G26A *sliP* variant resulted in neutrophil infections where significantly less deacetylase activity was detected than when the strain possessed complemented wild-type *sliP* (Figure 4A). To support these results, we also quantified NAD^+^ levels in similarly infected neutrophils. As expected, infected neutrophils possessing higher deacetylase activity due to the presence of wild-type SliP also exhibited significantly reduced NAD^+^ levels. In contrast, neutrophils infected with Δ*sliP* or the mutant complemented with the G26A variant exhibited significantly higher levels of intracellular NAD^+^ when compared to wild-type infected neutrophils (Figure 4B). No changes to NADH levels occurred under any condition tested (Figure 4B). These findings indicate SliP is translocated into neutrophils and the enzymatic activity of SliP promotes deacetylase activity in neutrophils, which likely impacts acetylation of various host proteins.

**Figure 4.**
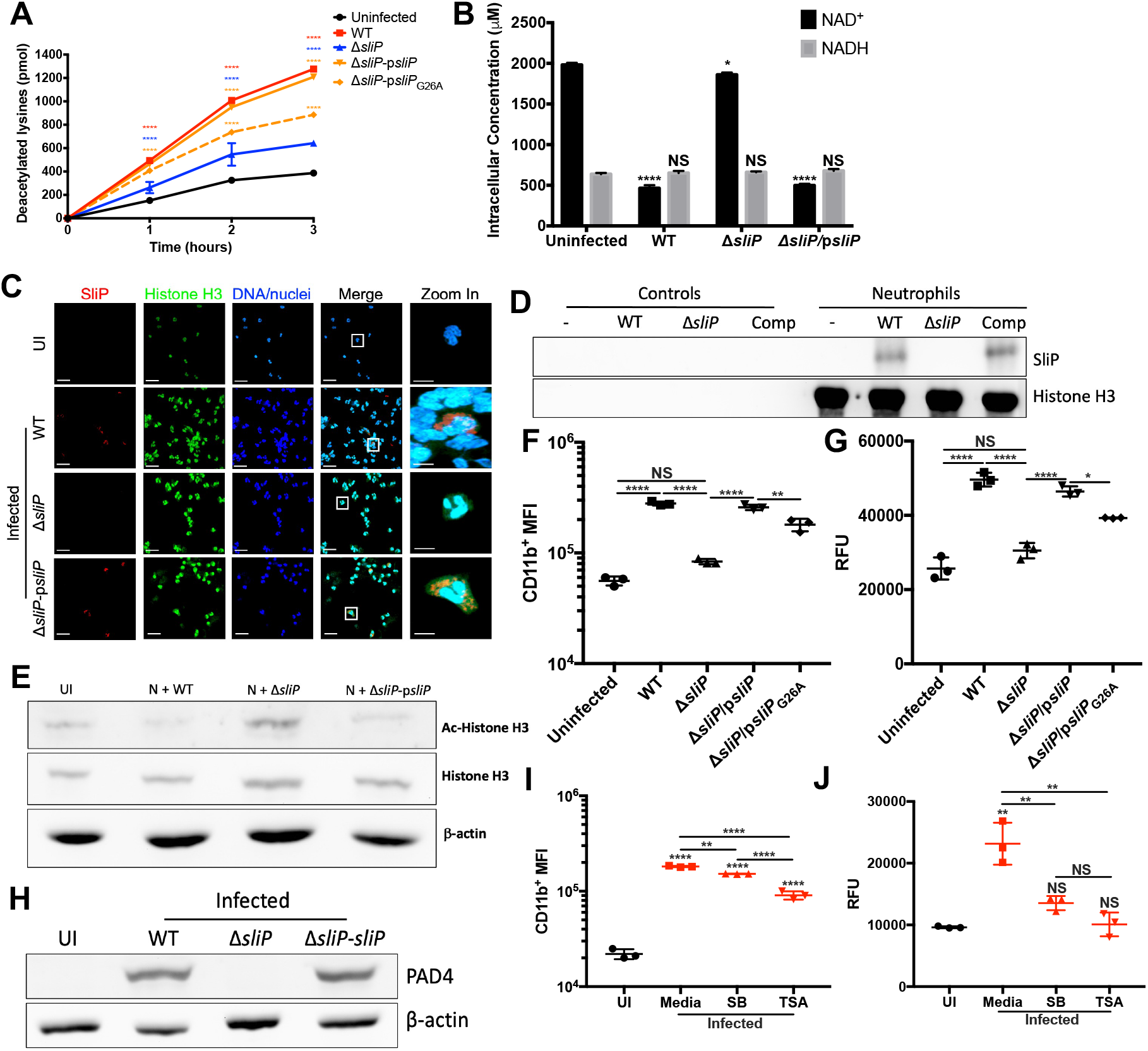
*C. jejuni* neutrophil activation and NET extrusion is dependent on the ability of SliP to bind and deacetylate histone H3. (A) Intracellular lysine deacetylation within neutrophils appears to be driven by SliP. Neutrophils were infected at an MOI of 10 and incubated under microaerobic conditions for up to three hours at 37°C. (B) Intracellular NAD^+^ and NADH concentrations within the neutrophils after being infected at an MOI of 10 for three hours at 37°C. (C) Fluorescent microscopy of secreted *C. jejuni* SliP co-localizing with histones during neutrophil infection. Uninfected (UI), wild-type (WT), Δ*sliP*, and *sliP* complemented (Δ*sliP*-p*sliP*) infected neutrophils were stained for nuclei/DNA (blue), histone H3 (green), and SliP (red) to observe intracellular SliP co-localizing with cytoplasmic histone H3. Scale bars are 20 μm for zoomed out images and 5 μm for zoomed in images. Images presented are composites of 3D reconstructions of z-stacked images. (D) Histone H3 co-immunoprecipitation of *C. jejuni* infected neutrophils. Neutrophils were incubated with *C. jejuni* at an MOI of 10 for three hours under microaerobic conditions. After incubation, cells were cross-linked with formalin and histone H3 was pulled down using an anti-histone H3 antibody. Presence of SliP within the histone H3 complex was determined through immunoblotting. (E) Immunoblot of SliP-dependent neutrophil histone H3 deacetylation after bacterial infection. (F) SliP-dependent *C. jejuni* induction of neutrophil activation through flow cytometry analysis of CD11b expression. (G) SliP-dependent *C. jejuni* induction of NETs through an extracellular DNA SYTOX assay and (H) PAD4 expression in neutrophils infected with *C. jejuni* expressing sliP. (I) Inhibition of SliP-dependent neutrophil activation and (J) NET generation using sodium butyrate (SB) and trichostatin A (TSA). Multiple comparison testing was performed using ANOVA with Tukey’s *post-hoc* test. \**p* < .05; \*\**p* <.01; \*\*\**p* < .001; \*\*\*\**p* < .0001

### SliP translocates to the cytoplasm during infection of neutrophils by *C. jejuni* and directly associates with histone H3

Following intracellular translocation, we hypothesized a likely target of SliP-dependent deacetylation were neutrophil histones. The rationale was based on observations that histone H3 is a frequent target of post-translational regulation and histone H3 is involved in several neutrophil behaviors, including PAD4-dependent hypercitrullination of deacetylated histone H3 prior to NET production. (Kenny et al., 2017). As expected, we observed colocalization of SliP with histone H3 under non-NETosing conditions. Interestingly, we observe this colocalization outside the nucleus of infected neutrophils, potentially due to chromatin decondensation following deacetylation (Figure 4C; Figure S6).

Because we were able to demonstrate colocalization of SliP with neutrophil cytoplasm and histone H3 during infection, we next sought to determine whether SliP binds neutrophil histone H3 during *C. jejuni* infection and promotes its deacetylation. To accomplish this, we immunoprecipitated histone H3 from primary human neutrophils that were formalin-fixed following infection with either wild-type *C. jejuni, ΔsliP*, or Δ*sliP*-p*sliP*. After purification, we found SliP was present in protein complexes containing histone H3 during neutrophil infection with either wild-type *C. jejuni* or Δ*sliP*-p*sliP* (Figure 4D). As expected, we did not detect SliP associating with histone H3 in Δ*sliP*-infected neutrophils (Figure 4D). To eliminate non-specific protein crosslinking as a potential explanation, we probed for cytoplasmic lipocalin-2 (Lcn2) in those same immunoprecipitated histone H3 complexes but were unable to detect Lcn2 (Figure S7A). This suggests SliP association with histone H3 was not due to non-specific crosslinking of proteins to neutrophil histones. As we observed SliP association with histone H3, we next sought to investigate whether SliP affects the acetylation of histone H3 during infection. In neutrophils infected with wild-type *C. jejuni*, histone H3 acetylation was significantly reduced by 9.57-fold when compared to the amount of histone acetylation present in uninfected neutrophils. Similar results were also observed for neutrophils infected with the complemented mutant (8.45-fold reduction of uninfected neutrophil levels). In contrast, Δ*sliP*-infected neutrophils displayed histone H3 acetylation intensities similar to those of uninfected neutrophils (1.12-fold reduction of uninfected result) (Figure 4E). To determine whether these deacetylation results may be due to the SliP-dependent recruitment of host sirtuins, we examined for the predominant human sirtuin, sirtuin-2 (Sirt2), in immunoprecipitated histone H3 complexes. Human Sirt2 was not detected in these complexes, which is a notable difference from other intracellular pathogens where the bacteria promote the recruitment of host Sirt2 to induce histone deacetylation (Figure S7A). To eliminate other host-derived factors that post-translationally modify histones, we examined the expression of all known histone H3-targeting HDACs and HATs during infection with *C. jejuni* and found none of these genes were significantly impacted (Figure S7B). Furthermore, we did not observe significant increases in Sirt2 abundance in infected neutrophils (Figure S7C). We therefore concluded SliP translocates to the cytoplasm of infected neutrophils and binds to histone H3 to directly deacetylate the protein rather than affecting the production or localization of a host-derived enzyme.

### SliP promotes neutrophil activation and extracellular trap formation by *C. jejuni* through its deacetylase activity

As host HDACs and sirtuins are necessary for neutrophil activation, NET production, and intracellular infection by other bacteria, we hypothesized the SliP may impact neutrophil behaviors through the above effects due to histone deacetylation. To assess this, we used wild-type *C. jejuni, ΔsliP, ΔsliP*-p*sliP*, and Δ*sliP*-p*sliP*_G26A_ to infect primary human neutrophils. Using CD11b^+^ as an output of neutrophil activation, wild-type infected neutrophils displayed a 5-fold increase in the number of CD11b^+^ cells as previously described (Callahan et al., 2020) while Δ*sliP*-infected neutrophils exhibited only a 1.49-fold increase when compared to uninfected neutrophils. Infection of neutrophils with Δ*sliP*-p*sliP* resulted in a 4.61-fold increase in the number of CD11b^+^ neutrophils while infection with Δ*sliP*-p*sliP*_G26A_ led to an intermediate 3.22-fold increase when compared to the uninfected neutrophils (Figure 4F). Furthermore, when NET production was evaluated, infection with wild-type *C. jejuni* led to a 93% increase in NET-associated DNA when compared to uninfected neutrophils using fluorescent staining of extracellular DNA (Callahan et al., 2020). Similar to our activation results, Δ*sliP*-infected neutrophils exhibited a 18.72% increase in extracellular DNA when compared to uninfected neutrophils. When *sliP* was complemented, extracellular DNA increased by 80.67% while infection with *sliP*_G26A_ variant-infected neutrophils exhibited a 52.91% increase in extracellular DNA when compared to uninfected neutrophils (Figure 4G). To support these observations, we also examined for SliP-dependent accumulation of PAD4 in *C. jejuni*-infected human neutrophils and obtained fold-changes relative to uninfected neutrophils of 12.95, 1.03, and 12.23 for wild-type, Δ*sliP*, and Δ*sliP*-p*sliP*, respectively (Figure 4H). To determine whether other bacterial deacetylases may be involved in neutrophil activation and NETosis, we examined the above phenotypes in a transposon mutant of the predicted intracellular sirtuin, *cobB*. In contrast to Δ*sliP* responses, all neutrophil activities elicited by *cobB::hawkeye* resembled those of wild-type infected neutrophils (Figure S8). As the presence of *sliP* is necessary for neutrophil activation and NETosis in response to *C. jejuni* infection, we concluded SliP plays a crucial role in modulating neutrophil responses to *C. jejuni* through its deacetylation of histone H3.

To determine if chemical inhibition of SliP deacetylase activity reduces neutrophil activation and NETosis, we preincubated neutrophils with SB and TSA at concentrations that inhibited SliP dependent deacetylation above. When pre-incubated neutrophils were infected with wild-type *C. jejuni*, we observed a significant 1.2 and 2.0-fold reduction in CD11b^+^ neutrophil populations when SB and TSA were added, respectively (Figure 4I). This reduced activation also resulted in a significant decrease in NETosis, with SB and TSA treatment leading to a 1.71-fold reduction and a 2.30-fold reduction in extracellular DNA, respectively (Figure 4J). Supporting this result, PAD4 levels were also significantly reduced in SB and TSA treated neutrophils during infection with wild-type *C. jejuni* (Figure S9). Furthermore, when SB and TSA treated neutrophils were infected with wild-type *C. jejuni*, histone H3 acetylation significantly increased (Figure S9). These results indicate SliP promotes neutrophil activation and NET induction through deacetylation of histone H3.

### SliP promotes inflammation and gastrointestinal disease during murine campylobacteriosis

To determine whether these SliP-dependent impacts on neutrophil activities affect host inflammation and gastrointestinal tissue damage during infection, we orally infected IL-10^-/-^ mice with either wild-type *C. jejuni* or Δ*sliP* and examined for systemic and tissue-level impacts on the host. Prior to orally infecting mice, we determined murine neutrophils behave similarly to human neutrophils when infected with *C. jejuni*, including activating similar numbers of neutrophils and inducing NETosis in those neutrophil populations (Figure S10). When IL-10^-/-^ mice were orally infected with wild-type *C. jejuni*, they exhibited significantly reduced percent weight change at days 7, 9, and 10 post-infection when compared to uninfected mice. In contrast, Δ*sliP*-infected mice gained weight similar to uninfected mice, which led to significantly higher weights when compared to wild-type infected mice at days 7 through 10 post-infection (Figure 5A). Importantly, Δ*sliP* was able to colonize mice, with similar fecal, colonic, and cecal loads as wild-type-infected mice (Figure 5B, C; Figure S11). In addition, we examined for bacterial dissemination from the gastrointestinal tract by determining the bacterial burden within the spleens of infected mice, finding Δ*sliP*-infected mice harbored 3.06-fold more bacteria than wild-type infected mice (Figure 5D). These results indicate SliP does not affect colonization of susceptible hosts and instead suggests neutrophil activation and NET induction could aid in restricting *C. jejuni* to the colon.

**Figure 5.**
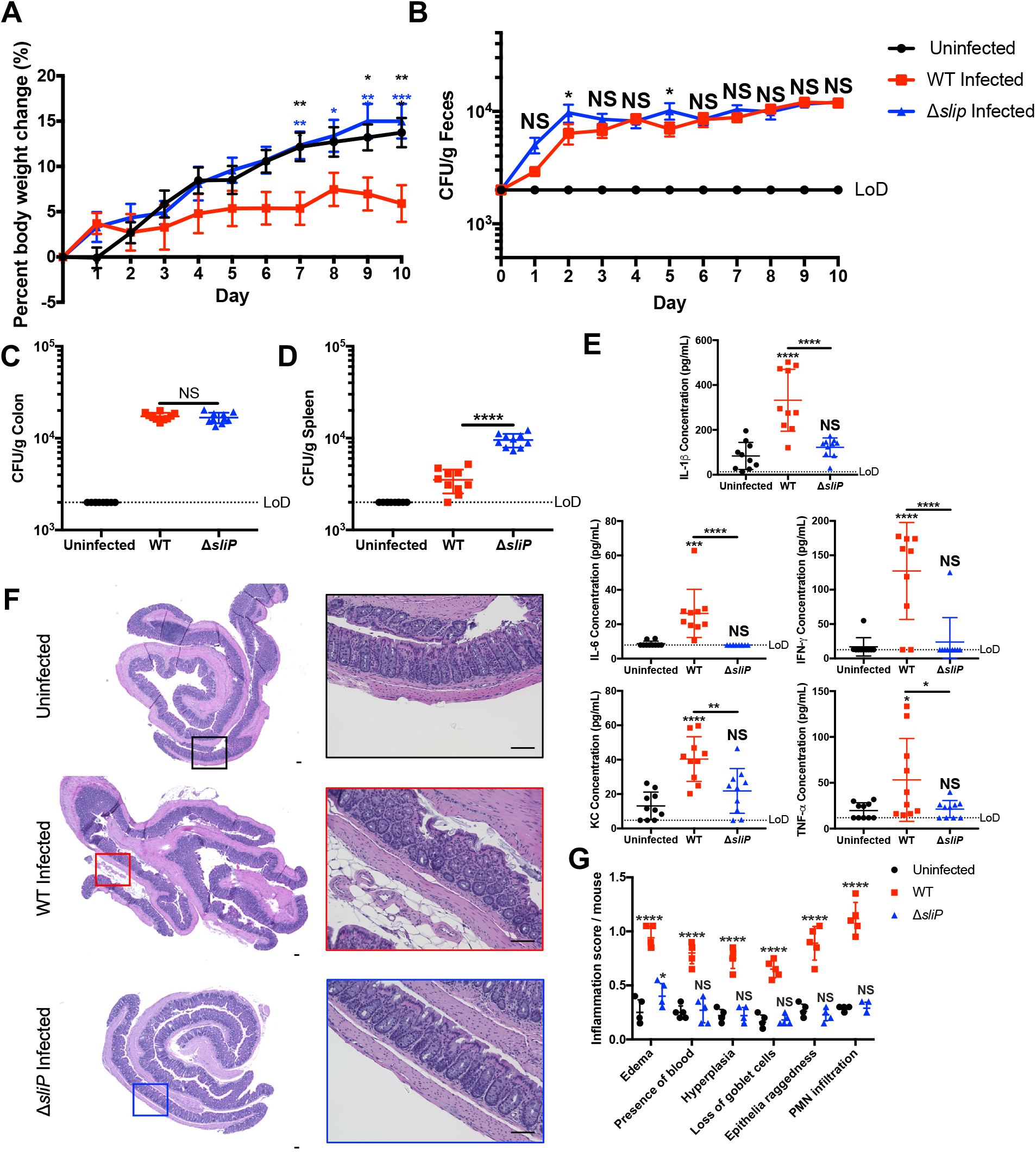
SliP promotes inflammation within the IL-10^-/-^ C57BL/6 mouse model of campylobacteriosis. (A) Percent body weight changes of mice uninfected or infected with wild-type (red) or Δ*sliP* (blue) *C. jejuni* over the ten-day infection study. Percent body weight changes were normalized to the weights of each mouse before savage treatment. (B) Fecal viable CFUs of each mouse over the ten-day infection study. Feces were collected each day and plated on *Campylobacter* specific media and incubated for two days under microaerobic conditions. (C) Viable CFUs from the colon and (D) spleen in *C. jejuni* infected mice ten days post-infection. (E) Serum cytokine expression levels for each mouse per treatment group. Levels of cytokines IFN-γ, KC, IL-1β, IL-6, and TNF-α were analyzed through flow cytometry-based beads and concentrations were determined through a standard curve for each protein. Multiple comparison testing was performed using ANOVA with Tukey’s *post-hoc* test. (F) H&E stained murine colons for each representative treatment group. Scale bars are 100 μm for zoomed out and in images for each treatment mouse group. (G) Inflammation scoring from the H&E stained colons per colon section. Analysis of inflammation scoring was performed using a nonparametric Mann-Whitney U test. \**p* < .05; \*\**p* < .01; \*\*\**p* < .001; \*\*\*\**p* < .0001

To determine whether SliP promotes inflammation and disease during infection, we first quantified the levels of several proinflammatory chemokines and cytokines within the serum of wild-type and Δ*sliP*-infected IL-10^-/-^ mice, including KC, IFN-γ, TNF-α, IL-1β, and IL-6. In uninfected mice, the concentration of KC was 13.13 ± 8.041 pg/mL, while concentrations in wild-type and Δ*sliP*-infected mice were 40.35 ± 12.99 pg/mL and 21.85 ± 12.99 pg/mL, respectively (Figure 5E). Similarly, the concentration of IFN-γ in uninfected mice was determined to be 16.95 ± 13.33 pg/mL, with wild-type and Δ*sliP*-infected mice exhibiting concentrations at 127.3 ± 70.49 pg/mL and 23.92 ± 35.55 pg/mL, respectively (Figure 5E). For TNF-α, uninfected mice concentrations were observed at 19.66 ± 8.53 pg/mL, while wild-type and Δ*sliP*-infected mice were at 53.13 ± 45.17 pg/mL and 21.31 ± 9.36 pg/mL, respectively (Figure 5E). For IL-1β concentrations, uninfected mice were observed at 83.86 ± 60.63 pg/mL, while wild-type and Δ*sliP*-infected mice were at 332 ± 138 pg/mL and 122.3 ± 41.81 pg/mL, respectively (Figure 5E). Finally, for IL-6 concentrations, uninfected mice were observed at 8.617 ± 1.535 pg/mL, while wild-type and Δ*sliP*-infected mice were at 26.33 ± 14 pg/mL and the limit of detection, respectively (Figure 5E). To investigate whether this SliP-dependent inflammatory response was also present in the colon, we examined leukocyte influx into the tissue, including the recruitment of neutrophils. In addition to promoting an immune response, we analyzed and scored colon pathology from wild-type and Δ*sliP*-infected mice, including the development of edema, blood in the tissue, epithelial raggedness, and hyperplasia (Lebeis et al., 2009). Importantly, when mice were infected with the Δ*sliP*, histopathological scores for the previously mentioned features were not significantly different from uninfected mice, aside from edema (Figure 5F, G). These results suggest despite colonizing mice at levels similar to wild-type and disseminating to extraintestinal sites, the absence of SliP leaves the bacterium less immunogenic and able to cause disease in a susceptible host.

### SliP associates with and promotes histone H3 deacetylation during murine infection

To connect the SliP-dependent activities we observed in primary human neutrophils to the pathological effects we observed in mice, we sought to demonstrate SliP association with histone H3 during infection. To initially examine this, colon sections from the above cohorts were analyzed by immunofluorescence microscopy. As expected, *C. jejuni* clusters were detected within the colons of wild-type and Δ*sliP* infected mice, but not within uninfected mice (Figure 6A). When histone H3 acetylation was examined within these clusters, histone acetylation co-localized in Δ*sliP* infected mice, however, this was absent in wild-type infected mice where histone acetylation was markedly reduced (Figure 6A). Further, myeloperoxidase (MPO), a leukocyte-derived antimicrobial protein, was also more abundant in wild-type-infected mice than uninfected and Δ*sliP*-infected mice (Figure 6A). To more specifically address whether SliP directly impacts the acetylation status of leukocyte histones during infection, colons from uninfected, wild-type, and Δ*sliP*-infected mice were harvested and CD45^+^ immune cells were purified. After purification, cellular proteins were cross-linked and histone H3 was immunoprecipitated as above. After purifying histone H3 complexes, we observed SliP was present in histone H3 complexes in only wild-type infected mice (Figure 6B, C). Furthermore, when the acetylation status of these complexes was examined, histone H3 acetylation was found to be reduced in wild-type-infected mice when compared to uninfected and Δ*sliP*-infected mice, both of which exhibited similar levels of acetylation (Figure 6D). As such, we have concluded SliP binds to histone H3 in murine immune cells and the resulting activation of those leukocytes is in part responsible for gastrointestinal disease. In the future, we will expand on these experiments to examine for SliP-dependent impacts to the acetylome of other cell types, including colonocytes and macrophages, and whether chemical inhibition of this activity reduces disease severity.

**Figure 6.**
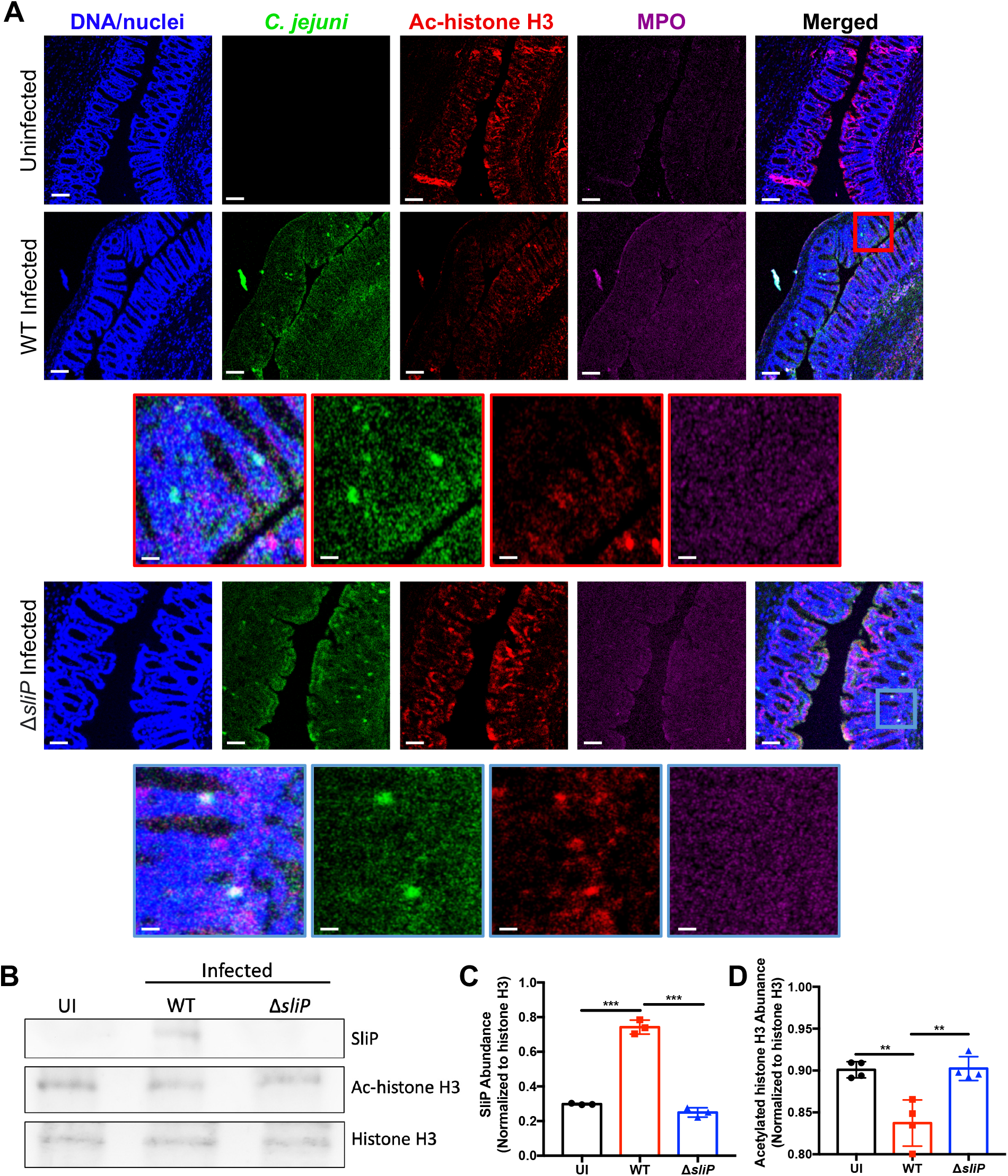
*In vivo* detection of SliP binding to and deacetylating immune cell histone H3. (A) *In vivo* fluorescent microscopy to determine the abundance of DNA/nuclei (blue), *C. jejuni* (green), acetylated histone H3 (red), and myeloperoxidase (MPO, purple) within uninfected, wild-type, and Δ*sliP* infected mouse colons. For each tissue section, the exposure for each channel remained constant. Images presented are individual channels and composites of z-stacked images. Scale bars are 100 μm for zoomed out images and 10 μm for zoomed in WT (red) and Δ*sliP* (blue) images. (B) Histone H3 co-immunoprecipitation from CD45^+^ immune cells purified from uninfected, wild-type, and Δ*sliP* infected mouse colons. Presence of SliP and the acetylation of histone H3 within the histone H3 complex was determined through immunoblotting. (C) Densitometry analysis of SliP abundance normalized to the abundance of histone H3 precipitated. (D) Densitometry analysis of histone H3 acetylation normalized to the abundance of histone H3 precipitated. Multiple comparison testing was performed using ANOVA with Tukey’s *post-hoc* test. \**p* < .05; \*\**p* < .01; \*\*\**p* < .001; \*\*\*\**p* < .0001

## Discussion

*Campylobacter* species are the leading cause of bacterial-derived gastroenteritis worldwide and have broad impacts on human health which range from acute intestinal inflammation to pediatric stunting (Chen et al., 2021). Despite these effects, knowledge relating to how infection with *Campylobacter* species results in these outcomes is limited. For example, because whole-genome analysis of *C. jejuni* isolates suggests the pathogen lacks many of the prototypical secretion systems and virulence factors often associated with pathogenesis, gastrointestinal disease is hypothesized to be due to dysregulated activation of the host immune response through a factor or process unique to *C. jejuni* (Ren et al., 2018). Since little work has been done to understand the host response to *C. jejuni* and its consequence on gastrointestinal health, our group previously examined and observed neutrophils are recruited to the colon of infected animals and that predominantly neutrophil-derived proteins are elevated in the feces of *C. jejuni*-infected patients. Because neutrophils play a key role in the development of several inflammatory diseases, some of which are related to the post-infectious disorders which potentially occur following *C. jejuni* infection, we worked to further characterize the neutrophil response to *C. jejuni*. For example, we recently demonstrated that *C. jejuni* is a potent activator of human neutrophils and stimulation leads to the production of NETs. Since neutrophil activities, including NETosis, can damage adjacent colonocytes and has been associated with numerous inflammatory disorders, we sought to identify unique factors of *C. jejuni* which promote neutrophil activation and NET extrusion.

In this work, we identified a unique *C. jejuni* effector which functions as a novel bacterial sirtuin and is released into neutrophils during intracellular infection. This protein, which we call SliP, is responsible for full neutrophil activation and NET production in response to *C. jejuni* infection. While several bacterial pathogens have been found to hijack host sirtuins to create an intracellular environment that promotes infection, an example of a bacterial-derived sirtuin being translocated into host cells to directly alter host responses has yet to be described. In addition to the enzymatic activity and proinflammatory functions of SliP, the gene is more frequently present in the genomes of human clinical isolates of *C. jejuni*. While the evolutionary rationale for this incidence is unknown, one explanation is that the role of SliP in neutrophil activation would lead strains which possess SliP to be more inflammatory, which would result in a detection bias since patients with severe disease are more likely to seek medical care where those strains are more likely to be isolated and sequenced. Beyond its role in *C. jejuni*, we found other bacterial pathogens encode DUF4917 family proteins; however, they have yet to be investigated. For example, the human pathogens *Legionella pneumophila* and *Yersinia enterocolitica* encode DUF4917 proteins with shared amino acid homology to *C. jejuni* SliP in addition to the multitude of other effector proteins these two pathogens possess. This observation suggests infection or disease caused by these pathogens may be similarly facilitated by the production of a bacterial-derived sirtuin.

Because we determined SliP acts as a canonical sirtuin *in vitro* and *in vivo*, we examined whether SliP inhibition could be achieved using the known deacetylase inhibitors, sodium butyrate and trichostatin A. Interestingly, sodium butyrate has been found to abrogate colonization of numerous gastrointestinal pathogens and reduce host inflammation, including those caused by *Salmonella* Typhimurium (Gupta et al., 2020; Rivera-Chávez et al., 2016), *Escherichia coli* (Xiong et al., 2016), *Citrobacter rodentium* (Ahmed et al., 2021; Jiminez et al., 2017), and *Staphylococcus aureus* (Park et al., 2019; Traisaeng et al., 2019). While the underlying molecular mechanism of this attenuation remains unclear, it may occur through inhibition of host-derived histone deacetylases (HDACs), which has been shown to have anti-inflammatory effects in other disorders, including inflammatory bowel disease. In addition, our results suggest sodium butyrate may reduce inflammation during infection by inhibiting the activities of bacterial-derived sirtuins that would otherwise alter the intracellular environment of the host to a more proinflammatory state.

Prior to identifying targets of SliP, we first examined whether SliP is secreted both *in vitro* and *in vivo*. As we found lysine deacetylation and NAD^+^ levels during *C. jejuni* infection of neutrophils were SliP dependent and that SliP is actively secreted into host cells, we hypothesized SliP targets host proteins to promote disease. Because HDAC upregulation and the resulting histone deacetylation promote NETosis and host inflammation, we initially focused on SliP-dependent impacts to the acetylation status of host histones. As predicted, we found SliP colocalizes and binds to histone H3 during neutrophil infection and that the presence of SliP leads to decreased histone acetylation. As we observed colocalization of SliP and histone H3 occurring outside the neutrophil nucleus, we hypothesize that this could be due to PAD4 dependent chromatin decondensation. Alternatively, SliP could be binding to histone H3 synthesized within the cytoplasm before shuttling into the nucleus for chromatin binding. We further demonstrated *C. jejuni*-induced neutrophil activation and NETosis is SliP-dependent, which suggests direct bacterial-mediated modification of host histones affect inflammatory responses and cell death. While we showed histone H3 becomes deacetylated in a SliP-dependent manner, other PTMs could occur after deacetylation, including lysine methylation, ubiquitination, and SUMOylation (Bernstein et al., 2002; Ryu and Hochstrasser, 2021; Zencheck et al., 2012; Zhang et al., 2017). Further supporting the role of SliP-dependent deacetylation in neutrophil activation, we found inhibition of SliP using the HDAC inhibitors discussed above reduced neutrophil activation and NET formation during *C. jejuni* infection. While it has been shown that *C. jejuni* can sense sodium butyrate through sensor phosphatase activity (Goodman et al., 2020), no research has been conducted on the ability of butyrate to influence host responses or pathogenesis during campylobacteriosis. Supporting the hypothesis that butyrate might reduce disease severity by affecting the host during infection, previous work found butyrate inhibits proinflammatory responses from neutrophils through downregulation of numerous antimicrobial proteins during mucosal inflammation models (Li et al., 2021a). Similarly, TSA has been shown to inhibit neutrophil proinflammatory responses; however, like butyrate, no research has been conducted to investigate the ability of TSA to reduce inflammation or disease during *C. jejuni* infection. As a result, the *C. jejuni* field is well-positioned to investigate the role of these inhibitors on campylobacteriosis disease severity and whether their abundance affects either bacterial and/or host-derived factors and processes.

Despite the ability of the *sliP* mutant to colonize and be shed in the feces similar to mice infected with wild-type bacteria, the bacterial burden within the spleen of *sliP* mutant-infected mice was significantly increased. We hypothesize this observation is similar to what was observed in a previous study where NET production was shown to be involved in limiting bacterial dissemination to the spleen during *Citrobacter rodentium* infection of mice (Saha et al., 2019). Furthermore, we observed *sliP* mutant infected mice have reduced levels of proinflammatory cytokines KC, TNF-α, IL-1β, and IL-6 compared to wild-type infected mice. We postulate either i) proinflammatory cytokines could be released by activated neutrophils during infection (Gideon et al., 2019; Tecchio et al., 2014), ii) tissue resident macrophages could detect citrullinated histone H3, resulting in upregulation of proinflammatory cytokines (Li et al., 2021b; Monteith et al., 2021; Tsourouktsoglou et al., 2020), or iii) influx of other leukocytes could be responsible for the increases of proinflammatory cytokines (Mangold et al., 2019; Parackova et al., 2020). Finally, we found murine inflammation and tissue pathology is dependent on the presence of SliP in *C. jejuni*. As neutrophil activation and NETosis has been implicated in numerous inflammatory diseases, whether autoimmune or microbially driven, we hypothesize SliP-dependent neutrophil responses drive campylobacteriosis disease manifestation.

With the characterization of a new class of bacterial effectors, sirtuins, the role of deacetylation on the host-microbe paradigm can be further understood. While microbial short-chain fatty acids have been implicated to abrogate bacterial-mediated gastroenteritis, future research can be focused on the ability to facilitate these interactions. By further understanding the unique mechanisms *C. jejuni* employs to promote infection and inflammation, novel therapeutics can be developed for treating acute and chronic outcomes of infection.

## Methods

### Abundance of SliP within environmental and clinical isolates

Environmental and clinical isolates of *C. jejuni* were isolated from a previous study (Kelley et al., 2020). Whole genomes were uploaded to KBase and their genomes were annotated to the 81-176 wild-type genome. After the genomes were annotated, amino acid homology was performed, with the minimum homology set to 75%. For statistical analysis, a two-sided Chi-square test was performed.

### Multiple sequence alignment of multi-organismal sirtuins

Bacterial and human sirtuins were identified as previously characterized in the literature. After the review of the literature, amino acids of each protein were acquired through NCBI. After each amino acid sequence for each protein was acquired, multiple sequence alignment was performed using T-Coffee (EMBL-EFI) with a ClustalW output. After the multiple sequence alignment was performed, alignments were aligned and colored using BoxShade. Conserved amino acids were identified using this coloring tool.

### Phylogenetic analysis of DUF4917 family proteins

To understand the homology of SliP with other members of the DUF4917 family, an amino acid BLAST was performed, excluding Campylobacter spp. DUF4917 proteins. From this BLAST search, amino acids were pulled from the top 14 percent homology organisms. The annotated amino acids were then aligned using multiple sequence alignment using T-Coffee (EMBL-EFI) with a ClustalW output. From this, a neighbor-joining tree without distance correction cladogram was produced.

### Bacterial cultures and culture conditions

*Campylobacter jejuni* 81-176 strain DRH212 was stored at −80°C in MH broth supplemented with 20% glycerol. In this study, non-polar deletions and plasmid-borne strains of *sliP* were constructed in wild-type *C. jejuni* DRH212 as previously described (Hendrixson and DiRita, 2003). All strains were routinely grown for 24 hours on selective media (10% sheep’s blood, 40 μg/ml cefoperazone, 100μg/ml cycloheximide, 10 μg/ml TMP and 100μg/ml vancomycin) before passing onto Mueller-Hinton agar containing 10% sheep’s blood and 10 ug/ml trimethoprim (TMP) at 37°C under microaerobic conditions (85% N2, 10% CO2 and 5% O2) for an additional 24 hours. *E. coli* expression strain C3013 containing pQE30-*sliP* and pQE-30-*sliP*_G26A_ were constructed as previously described (Kelley et al., 2021). *E. coli* strains were stored in Luria-Bertani (LB) broth supplemented with 20% glycerol at −80°C and grown shaking at 37°C in LB broth supplemented with ampicillin for purification as described below.

### Isolation of primary human neutrophils

Neutrophils were isolated as previously described (Callahan et al., 2020, 2021b). Briefly, 10 ml of blood was drawn from healthy donors into heparinized Vacutainer tubes and mixed 1:1 with sterile 1× PBS (approval UTK IRB-18-04604-XP) (Akhtar et al., 2010). After mixing, 10 ml of lymphocyte separation medium was underlaid. Following centrifugation at 1,400 rpm for 30 min, the top layers were aspirated off, leaving the red blood cell and neutrophil pellet. Pellets were resuspended in 20 ml Hanks’ balanced salt solution and 20 ml 3% dextran in 0.9% NaCl. After incubating at room temperature for 20 min, the upper layer was transferred to a new tube. Following centrifugation at 400*g* for 5 min and aspiration of the supernatant, the pellet was washed with 20 ml ice-cold 0.2% NaCl and 20 ml ice-cold 1.6% NaCl two times. Following the final aspiration, the neutrophil pellet was resuspended in 10 ml RPMI 1640. Neutrophil viability and counts were performed through Trypan blue stain.

### Protein induction and native purification of SliP

Protein induction and purification was performed as previously described (Johnson et al., 2016; Kelley et al., 2021). Briefly, *E. coli* strains C3013 encoding SliP, SliP_G26A_, or empty pQE-30 vector were grown overnight in LB broth containing 100 μg/ml ampicillin at 37°C shaking. Cultures were then back-diluted by adding 1 mL overnight culture in 100 mL LB broth containing 100 μg/ml ampicillin. Culture was grown at 37°C shaking for two hours before spiking in 100 μM isopropyl-β-d-thiogalactopyranoside (IPTG) to induce protein expression. Cultures were allowed to incubate for an additional two hours at 37°C shaking. After incubation, cells were pelleted at 2,147 rcf for 10 min. After centrifugation, the pellets were resuspended in lysis buffer (50 mM NaH_2_PO_4_, 300 mM NaCl, 10 mM imidazole, pH 8.0) before sonication on ice (6 rounds of 15 s each at 45 A). Cell lysates were then pelleted at 2,147 rcf for 10 min before incubating the resultant supernatant with washed Ni-nitrilotriacetic acid (Ni-NTA) for one hour at 4°C rocking. After the incubation, the SliP resin was packed in a 20-ml chromatography column before it was washed three times with 4 ml wash buffer (50 mM NaH_2_PO_4_, 300 mM NaCl, 20 mM imidazole, pH 8.0) and eluted in three 750-μl fractions of elution buffer (50 mM NaH_2_PO_4_, 300 mM NaCl, 250 mM imidazole, pH 8.0). Each wash and elution fraction were diluted 1:1 with a Laemmli buffer containing β-mercaptoethanol and boiled for 10 minutes. Ten microliters of lysate samples were loaded onto a 12.5% SDS-PAGE gel and run for 1 hour at 150 V at room temperature and stained with Coomassie to ensure protein purification (37 kDa). Elution fractions were then pooled and dialyzed using a dialysis buffer (200mM NaCl, 2mM DTT, and 20mM HEPES buffer (pH 7.5) three times at 4°C spinning. After dialysis, protein was collected from the dialysis cassette using a syringe and stored at 4°C until future use.

### Detection of *in vitro* and *in vivo* lysine deacetylation

Quantification of *in vitro* and *in vivo* lysine deacetylation was performed using the BioVision InSitu HDAC Activity Fluorometric Assay kit (Cat. No. K339-100). For *in vitro* quantification, 10 ng purified SliP was incubated in the presence of HDAC substrate (acetylated lysine side chain) with 5 mM NAD^+^ and 20 μM ZnCl_2_ at 37°C. For *in vivo* quantification, 10^5^ neutrophils were incubated with 10^6^ *C. jejuni* to a final volume of 150 μl containing the HDAC substrate and the cells were infected as previously described. After the allotted amount of time, developer and HDAC assay buffer was added to 100 μl reaction mixture (with the other 50 μl being used for the NAD^+^ hydrolysis assay described below) and was incubated for 30 minutes at 37°C. After incubation, fluorescence was read using a BioTek Synergy microplate reader at 368/442 nm wavelength. A deacetylated substrate standard curve was additionally run and plotted with fluorescence. Once plotted, the amount of deacetylated lysine residues were calculated.

### Quantification of NAD^+^ and NADH within neutrophils and supernatants

The abundance of NAD+ and NADH in vitro and in vivo were performed using the Promega NAD/NADH-Glo™ Assay. All reagents used for this assay were brought to room temperature before use. For the *in vitro* NAD^+^ hydrolysis assay, 5 mM of pure NAD^+^ was added to the reaction as described above in the lysine deacetylation assay. After deacetylation, the NAD/NADH-Glo™ detection reagent was made as described by the manufacturer. In a white welled plate, 50 μl from the lysine deacetylation reaction mixture was combined with 50 μl detection reagent. The plate was incubated at 37°C for 30 minutes while shaking. After incubation, luminescence was read using a BioTek Synergy microplate reader. For quantification of intracellular NAD^+^ and NADH, neutrophils were infected with their respective strain for the allotted amount of time under microaerobic conditions at 37°C to a final volume of 50 μl. Neutrophils were then lysed by adding 50 μl RPMI 1640 containing 1% dodecyltrimethylammonium bromide (DTAB). To measure NAD^+^, 50 μl of the reaction mixture was allotted and mixed with 25 μl 0.4N hydrochloric acid and incubated at 60°C for 15 minutes. After the incubation, the samples were cooled at room temperature for 10 minutes. Then, 25 μl Trizma base was added to the reaction mixture. Finally, 100 μl of detection reagent was added, incubated as previously described and luminescence was read as previously described. For NADH quantification, 50 μl of the cell lysis solution was incubated at 60°C for 15 minutes. The samples were cooled at room temperature for 10 minutes. Then, 50 μl HCl/Trizma base solution was added to the reaction mixture. Finally, 100 μl of detection reagent was added, incubated as previously described and luminescence was read as previously described. Along with each experiment, a known concentration standard curve for NAD^+^ and NADH were constructed to calculate the concentration of each small molecule.

### SliP-CyaA neutrophil translocation assay

To determine if SliP is translocated into neutrophils, adenylate cyclase translational fusions were produced at the C terminus of *sliP, heuR*, and *ciaB* (Sory and Cornelis, 1994). For each infection, 10^6^ neutrophils were incubated with wild-type or Δ*flgE C. jejuni* strains harboring pECO102 encoding *sliP-cyaA, heuR-cyaA*, or *ciaB-cyaA* at an MOI of 10 for three hours under microaerobic conditions at 37°C. Additionally, phagocytosis was chemically blocked with 2 μM cytochalasin D (CD) for one hour before bacterial infection under microaerobic conditions at 37°C. Cells were lysed with 0.1M HCl and the direct ELISA was performed as according to the manufacturer protocol (Enzo).

### Analysis of *C. jejuni* NET induction by flow cytometry

Flow cytometry detection of *C. jejuni* induced NETs was performed as previously described (Callahan et al., 2020, 2021b). Briefly, *C. jejuni* strains were resuspended in RPMI 1640 and incubated with neutrophils for 3 hours under microaerophilic conditions at an equal multiplicities of infection (MOI) of 50:1 (bacteria: neutrophil). Following incubation, these reactions were aliquoted into wells of a 96-well plate and centrifuged at 400 x *g*. Pelleted cells were washed three times with sterile 1x PBS cells before incubation with Live/Dead Near-IR (NIR) stain for 25 min at room temperature. After incubation, cells were blocked with 1% goat serum for 10 min, then incubated with CD11b and MPO antibodies for 30 min (Zharkova et al., 2019). Cells were subsequently fixed in the fixation buffer for 10 min at room temperature and stored at 4°C until flow cytometry analysis. Samples were analyzed using an LSR II flow cytometer and data were processed using BD FlowJo software. Statistical analysis was performed using unpaired *t* tests and significance inferred at *p* < .05.

### Analysis of *C. jejuni* NET induction by SYTOX assay

SYTOX NET quantification was performed as previously described (Callahan et al., 2020, 2021b). Briefly, Following the *C. jejuni* NET induction described above, cells were added to wells of black 96-well plates and centrifuged at 400 x *g*. Supernatants were discarded and 1.0 μM SYTOX Green was added to each sample and incubated under microaerobic conditions at 37°C for 1 hr (Brinkmann et al., 2010). After incubation, cells were centrifuged at 400 x *g* and the supernatant was discarded. The cells were washed once with 1× PBS and then resuspended in 100 μl 1× PBS. SYTOX fluorescence was measured at 504/523 nm using a BioTek Synergy microplate reader. Statistical analysis was performed using unpaired *t* tests and significance inferred at *p* < .05.

### Western blot analysis of NET products

Detection of NET components via western blot was performed as previously described (Callahan et al., 2020). Briefly, NETs were induced as described above, cells from healthy volunteers were lysed with Laemmli buffer with β-mercaptoethanol and boiled for 10 min. Ten microlitres of whole-cell lysate samples were loaded onto a 12.5% SDS-PAGE gel and run for 1.25 hr at 140 V at room temperature. Separated proteins were transferred to nitrocellulose membranes (GE Healthcare, Cat. 10600011) for 1.5 hr at 0.25 A using a semi-dry transfer apparatus. Membranes were blocked in 5% milk in Tris-buffered saline (TBS)-T for 60 min shaking. To probe proteins of interest, 1:500 dilution rabbit anti-*C. jejuni* SliP (Cocalico Biologicals), 1:5000 dilution rabbit anti-human Lcn2 (Invitrogen, Cat. 702248), 1:1000 dilution rabbit anti-human peptidylarginine deiminases 4 (PAD4; Thermo, Cat. PA5-12236), 1:1000 dilution rabbit anti-human histone H3 (Active Motif, Cat. 61799), 1:1000 dilution rabbit anti-human acetyl histone H3 (Active Motif, Cat. 39040), and 1:500 dilution mouse anti-human β-actin (Cell Signaling, Cat. 4970) were incubated on individual membranes for 1 hr at room temperature with shaking. Membranes were subsequently washed three times with filtered TBS-T for 5 min each. Proteins of interest were detected using a 1:2000 dilution of appropriate horseradish peroxidase-conjugated secondary antibodies in 15 ml 5% milk in TBS-T and incubated for 45 min at room temperature with shaking. Membranes were then washed three times with filtered TBS-T, shaking for 5 min each. After the final wash, membranes were developed with a solution containing 5 ml peroxide and 5 ml luminol/enhancer solution (Thermo, Cat. 34580) and incubated for 5 min at room temperature with shaking. A *C. jejuni* negative control was also performed to ensure there was no cross-reactivity and no detection of proteins was noted. Chemiluminescent bands were imaged using the ChemiDoc-It Imager and densitometry measurements were made using NIH ImageJ software. Statistical analysis was performed using unpaired *t* tests and significance inferred at *p* < .05.

### RT-qPCR analysis of gene expression

For determining *C. jejuni sliP* expression during neutrophil infection, we used a standard phenol-chloroform RNA extraction as previously described (Kelley et al., 2021). Briefly, cells were centrifuged at 21,130 x G for 5 minutes. The supernatant was aspirated off and the cells were resuspended in 500 μl RiboZol RNA extraction reagent (VWR). Tubes were then vortexed and 100 μl chloroform was added to the tube. Tubes were then centrifuged at 21,130 x G for 15 minutes at 4°C. The aqueous phase was then pipette off and placed in a tube containing 250 μl isopropanol. Nucleic acids were pelleted through centrifugation at 21,130 x G for 15 minutes at 4°C. Nucleic acid pellets were washed in 70% ethanol and air dried at room temperature. Nucleic acid pellets were then resuspended in nuclease-free water and stored at −20°C. For extracting neutrophil RNA, RNA was extracted using a Qiagen RNeasy Kit per the manufacturer protocol. RNA samples were eluted from the kit columns using nuclease-free water and stored at −20°C. All RNA samples were then DNase treated, and pure RNA was clean and concentrated using a kit (Zymo). Each sample was tested for DNA contamination through standard PCR. cDNA was then produced from DNA-clear RNA using the BioRad iScript kit using the manufacturer protocol. Primers specific to *sliP, HDAC1, HDAC2, HDAC3, SIRT1, SIRT2, SIRT6, SIRT7, GCN5, PCAF, P300, CBP, SRC1, ACTR, TAF11250, TFIIIC90, GAPDH, and rpoA* were designed using PrimerQuest (IDT) (Table S1). The abundance of each gene expression was determined using the iTaq Universal SYBR green master mix (BioRad). Threshold values (*C_T_*) were used to calculate the ΔΔ*C_T_* using the *GAPDH* and *rpoA* as the internal control for human and *C. jejuni* genes, respectively. For *sliP* gene expression, abundance was normalized to *C. jejuni* in media alone. For neutrophil gene expression, abundance was normalized to neutrophils in media alone.

### *In vitro* immunofluorescence microscopy

Cocultures were completed in wells containing sterile 12 mm poly-L-lysine coated coverslip (Corning, Corning NY) and cells were fixed in a solution of 2% paraformaldehyde, 2.5% glutaraldehyde in 0.1M sodium cacodylate buffer (pH 7.4) (Electron Microscopy Sciences, Hatfield, PA). Coverslips were washed thrice with tris buffered saline (TBS, pH 7.4) before permeabilizing with 0.25% Triton-X 100. Samples were then incubated in blocking buffer (TBS with 10% bovine serum albumin) for 120 minutes followed by overnight incubation at 4°C with a 1/200 dilution of the poly-clonal rabbit sera and antibodies for histone H3 (final concentration 2.5 μg/mL, Sigma SAB4200651) in blocking buffer. Samples were washed in blocking buffer thrice and stained with 1:1000 dilution of Alpaca anti-Rabbit IgG Nano (VHH) Recombinant Secondary Antibody conjugated with Alexa Fluor™ 647 (Invitrogen SA5-10327), Goat anti-Mouse IgG conjugated with Alexa Fluor™ Plus 488 (5 μg/mL final concentration, Invitrogen A32723), and 10 μM Hoechst 33342 for 120 minutes at room temperature. Coverslips were washed twice in blocking buffer and twice in TBS before mounting with ProLong™ Glass Antifade Mountant (Invitrogen P36982). Samples were visualized with a Zeiss LSM 710 META inverted laser scanning confocal microscope in part through the use of the Vanderbilt Cell Imaging Shared Resource. Presented images are 3D reconstructions of z-stacked images.

### Co-immunoprecipitation of histone H3 and SliP

Neutrophils were purified and infected with *C. jejuni* wild-type, Δ*sliP*, and complement strains as previously described. After non-NETosing condition incubation, cells were fixed and cross-linked by the addition of paraformaldehyde to a final concentration of 4% and incubated for 10 minutes at room temperature. After cross-linking, glycine was added to a final concentration of 125 mM and incubated for 5 minutes, shaking. The cells were then washed three times with ice cold 1x PBS. After the final wash, the cells were lysed using 750 μl RIPA buffer (150 mM NaCl, 1% Nonidet P-40, 0.5% deoxycholate, 0.1% sodium dodecyl sulfate, and 50 mM Tris (pH 7.4)). After lysis of the cells, the samples were centrifuged for 10 min at 4°C, 14,000 x g. After centrifugation, the supernatant was pulled off and placed in a sterile 1.5 mL Eppendorf tube. Next, the histone H3 antibody was added to the supernatant at a dilution of 1:1000. The supernatant-antibody solution was incubated overnight at 4°C rocking. After the incubation, 10 μl protein A beads were added to each sample and incubated at 4°C for 3 hours rocking. After incubation, the tubes were placed on a magnet to pull down beads conjugated to the antibody-protein complexes. The conjugated beads were washed three times with RIPA buffer, using a magnet at 4°C each time. After the last wash, the proteins were eluted by adding Laemmli buffer with β-mercaptoethanol and boiled for 10 minutes. After the boiling, the tubes were placed on a magnet to discard the protein A agarose beads. Eluted proteins in Laemmli buffer were stored at −20°C until western blot analysis as previously described.

### Infection of IL-10^-/-^ mice with *C. jejuni*

All animal protocols were approved by the Institutional Animal Care and Use Committee at the University of Tennessee - Knoxville (UTK IACUC protocol #2885). As previously described, *C. jejuni* DRH212 and Δ*sliP* were grown on *Campylobacter*-specific medium and Gram stained to ensure culture purity prior to inoculation (Shank et al., 2018). This culture was streaked on *Campylobacter*-selective media and incubated at 37°C under microaerophilic conditions for 48 hr. Suspensions of *C. jejuni* 81–176 were made in sterile PBS and diluted to an OD600 of 10. Five female specific pathogen free 8- to 12- week old IL-10^-/-^ C57BL/6 mice were gavaged with a single dose of 10^10^ CFUs per mouse of *C. jejuni* wild-type strain 81-176 or the Δ*sliP* mutant. Additionally, five female mice were mock infected with sterile 1× PBS. Each day for 10 days, approximately 20 mg of feces from one sample from each animal was immediately weighed out and diluted 1:100 in PBS. The remaining samples were immediately frozen at −80°C. Diluted samples were further serially diluted in PBS to 10^−8^, and 100 μl of each dilution was plated on *Campylobacter*-specific medium. The plates were then incubated at 37°C under microaerophilic conditions for 48 hours before *Campylobacter* loads were determined. After 10 days post-infection, blood was collected by cardiac puncture and mice were euthanized as previously described. From the blood, serum was collected and frozen at −20°C for further analyses. Ceca, colons, and spleens were excised from each mouse and placed in sterile 1X PBS. Colons were additionally rolled in Swiss Rolls and placed in formalin solution before being fixed in paraffin. To determine bacterial load in these tissues, ground tissue was weighed and diluted 1:100 in PBS. Diluted samples were further serially diluted in PBS to 10^−8^, and 100 μl of each dilution was plated on *Campylobacter*-specific medium.

### Histological analysis of murine intestinal tissue during *C. jejuni* infection

Pathology scoring for intestinal tissue was performed as previously described (Callahan et al., 2020). The terminal 1 cm of the murine colon was removed following sacrifice, and the lumen was washed five times with 1 ml of sterile, cold 1× PBS. Tissue was placed in 10% buffered formalin and fixed for 4 hours at room temperature. Fixed tissue was embedded in formalin, and 4 μm sections were stained with haematoxylin and eosin (H&E). Stained slides were visualized through bright-field microscopy, and representative images were presented. Inflammation scoring was performed as described in a blind manner, noting oedema, presence of blood, hyperplasia, loss of goblet cells, epithelia raggedness and neutrophil infiltration (Lebeis et al., 2009). Statistical analysis was performed using a nonparametric Mann-Whitney U test.

### Quantification of proinflammatory cytokines from murine sera

Murine cytokines from sera were quantified using the BioLegend LEGENDplex Anti-Virus response panel in a V-bottom plate per the manufacturer guidelines. Briefly, sera was plated, alongside a standard curve of known concentrations for various cytokines. Assay buffers were added to each well, along with mixed beads that had been sonicated in water baths and vortexed to ensure homogeneity. The plate was sealed, wrapped in aluminum foil, and shook on a plate shaker at 800 rpm for two hours at room temperature. After incubation, the plate was centrifuged at 250 X G for 5 minutes using a swinging basket. After centrifugation, the plate seal was removed, and the supernatant was immediately flicked off into a bucket. The wells were then washed using the manufacturer wash buffer twice before the detection antibody was added to each well. The plate was sealed and covered in aluminum foil and shook on a plate shaker at 800 rpm for one hour at room temperature. After incubation, a streptavidin-phycoerythrin solution was added to the wells and the plate was sealed and wrapped in aluminum foil, where it incubated on a plate shaker at 800 rpm for 30 minutes at room temperature. After incubation, the plate was centrifuged and washed as previously described. Once the preparation was complete, the samples were resuspended in wash buffer until flow cytometry analysis. Samples were analyzed using an LSR II flow cytometer and data were processed using the BioLegend LEGENDplex data analysis software. Cytokine concentrations were determined due to the standard curve generated by standard bead concentrations.

### *In vivo* fluorescence microscopy of mouse tissue samples

Parrafin embedded mouse tissues were stained for fluorescence microscopy as previously described (Brinkmann et al., 2016; Callahan et al., 2020; Doster et al., 2018). Briefly, slides containing slices of paraffin embedded tissues were digested in three rounds of xylene washes for five minutes, followed by a final wash in xylene for 10 minutes at room temperature. Afterwards, slides were washed with two rounds of 5-minute washes with 100%, 95%, 70%, 50% ethanol each at room temperature. After the final ethanol wash, slides were rehydrated in deionized water by incubating for 5 minutes twice at room temperature. After rehydration, slides were incubated in R universal Epitope Recovery Buffer (Electron Microscopy Sciences) at 50°C for 90 minutes. After incubation, slides were then washed with deionized water three times for 5 minutes each. The slides were then washed with TRIS-buffered saline (TBS, pH 7.4) three times for 5 minutes each at room temperature. Samples were then permeabilized for 5 minutes with 0.5% Triton X100 in TBS at room temperature followed by three washes with TBS for 5 minutes each at room temperature. Samples were then blocked with TBS with 10% BSA for 30 minutes at room temperature. After blocking, samples were then incubated with 1:20 dilution of histone H3, 1:20 acetyl-histone H3, 1:50 Campylobacter, and 1:100 MPO antibodies overnight at room temperature covered in the dark. After overnight incubation, slides were washed three times with TBS for 5 minutes each at room temperature. After the final wash, the samples were stained with 5 μM Hoechst 33342 for 30 minutes at room temperature. After the incubation, slides were washed three times with TBS for 5 minutes each at room temperature. After the final wash, the slides were air dried and coverslipped using Mowiol. Tissues were visualized with a Nikon E600 Eclipse at the Advanced Microscopy and Imaging Center at the University of Tennessee. Images presented are composites of z-stacked images.

### Purification of immune cells from murine colon tissues for co-immunoprecipitation

CD45^+^ immune cells were purified from the colons of uninfected, wild-type, and Δ*sliP* infected mice per manufacturer guidelines (MojoSort Biolegend). After purification of CD45^+^ cells, cells were incubated in formalin to cross-link protein complexes at 4°C. The histone H3 co-immunoprecipitation was performed as previously described, with samples stored at −20°C until western blot analysis was performed.

## Acknowledgments

Support for this project was provided by the University of Tennessee as start-up funds to J.G.J. The authors would like to thank Dr. David Hendrixson at UT Southwestern for supplying the *C. jejuni* 81-176 *flgE* mutant. The authors would additionally like to thank Dr. Chris Price and Dr. Yousef Abu Kwaik at the University of Louisville for supplying the *ankB-cyaA* adenylate cyclase vector plasmid.

## Author Contributions

Conceptualization, S.M.C. and J.G.J.; Methodology, S.M.C. and J.G.J.; Investigation, S.M.C., T.J.H, R.S.D., and J.G.J.; Writing – Original Draft, S.M.C. and J.G.J.; Writing – Review & Editing, S.M.C., T.J.H., R.S.D., C.B.P., M.E.W., J.A.G., J.G.J..; Funding Acquisition, R.S.D., J.A.G. and J.G.J.; Resources, J.A.G. and J.G.J.; Supervision, J.G.J.

