## Supplemental Movie for "Battle for the histones: a secreted bacterial sirtuin from *Campylobacter jejuni* activates neutrophils and induces inflammation during infection"

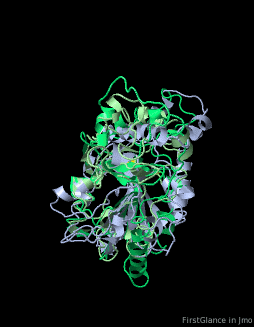

**Supplemental Movie 1**. Rotating predicted model according to secondary structure homology. Yellow dots depict the position of the G26 amino acid for SliP in the predicted Rossman fold domain. Overlapping structures were generated and the movie was produced in Jmol software.
