## Supplemental Figures for "Battle for the histones: a secreted bacterial sirtuin from *Campylobacter jejuni* activates neutrophils and induces inflammation during infection"

**Supplementary Information**

**Table S1**: Primers used in the study

**Primers used for making the bacterial strains** (5'-3'):

| | |
|---|---|
| *sliP* F | GTATATTCGCGTATGGAGTGG |
| *sliP* R | GGATTACCTGCTCCATTTGCT |
| SOE *sliP* F | ATGGAAAATAAACTATTAGAAGGATCCAAAAGTGTAAAAATTTGGTAA |
| SOE *sliP* R | TTACCAAATTTTTACACTTTTGGATCCTTCAAATAGTTTATTTTCCAT |
| *sliP* complement F | GGATCCATGGAAAATAAACTATTTGA |
| complement *sliP* complement R | GGATCCTCCTTGTTTTTACCAAATT |
| JJ13 | TGCTCTAGATTTATGATATAGTCGATA |
| *sliP* G26A F | GACACCTTCTTTTAGGAAATGCTTTTAGTATGGCTTATGAT |

| | |
|---|---|
| *sliP* G26A R | ATCATAAGCCATACTAAAAGCATTTCCTAAAAGAAGGTG |
| *sliP-cyaA* SOE F | TATAAAAGTGTAAAAATTTGGGCTGCTCAGCAATCGCATCAG |
| *sliP-cyaA* SOE R | GCTGAGCAGCCCAAATTTTTACACTTTTATA |
| *heuR-cyaA* SOE F | TATCTGAAGAAATTTAAAAAAGCTGCTCAGCAATCGCATCAGGC |
| *heuR-cyaA* SOE R | CGATTGCTGAGCAGCTTTTTTAAATTTCTTCAGATA |
| *heuR* complement F | GGATCCATGGATGAGGGACAAAAACAA |
| *ciaB-cyaA* SOE F | TTTGAAAGATATAAGAAAAAAGCTGCTCAGCAATCGCATCAGGC |
| *ciaB-cyaA* SOE R | GCTGAGCAGCTTTTTTCTTATATCTTTCAAA |
| *ciaB* complement F | GGATCCATGAATAATTTTAAAGAAATAGCTAAATTGG |

| | |
|---|---|
| *cyaA* complement R | GGATCCTGTCATAGCCGGAATCCTGGC |

**Primers used for RT-qPCR experiments** (5'-3'):

| | |
|---|---|
| *sliP* qPCR F | TGGTGGTAATGATGGAGAATATGT |
| *sliP* qPCR R | TTCTGATGTGCGTTCCGATAC |
| *HDAC1* F | GGAAGAGGAGTGAGCATTAGAG |
| *HDAC1* R | CTACCTTGGGATTGGGTTAGAG |
| *HDAC2* F | GGAGGGTCTCTTGTCTGTATTG |
| *HDAC2* R | TGGGTCATGCGGATTCTATG |
| *HDAC3* F | TGCATTGTGCTCCAGGTAATA |
| *HDAC3* R | CCTTCCACCACCAACCTAAA |
| *SIRT1* F | GCCCGGTCAGGTTTCTTATT |
| *SIRT1* R | CCTCCCAAAGTGCTAGGATTAC |

| | |
|---|---|
| *SIRT2* F | GAAAGGACCCTGGCTACTAAAG |
| *SIRT2* R | TCCCAATGTGCTGGGATTAC |
| *SIRT6* F | GTGGACATCGCCTTCTCTAATC |
| *SIRT6* R | GACCAGGAGAAACAGGAACAA |
| *SIRT7* F | CTGTTACTCTCACTCGGCTTTC |
| *SIRT7* R | CGTCATCACACTTCCCATGTAG |
| *Gcn5* F | CTAAAGGAGGGTGTGAGTGAAG |
| *Gcn5* R | AGTAGCTAGAGAGAAGAGGAAGG |
| *PCAF* F | GTGGGACATCCTTGACACTAAT |
| *PCAF* R | CATCCAGAGGAACAGGAGAAAG |
| *P300* F | CGGCCAGAGGTACCATTATATC |
| *P300* R | GAGGTGATGTGCCTCCAATAA |
| *CBP* F | CTCCCAGGTTCAAGCTATTCTC |

| | |
|---|---|
| *CBP* R | CGAAACCCTGTCTCCACTAAA |
| *SRC-1* F | GGAGGACGAAGACTGAACATAAA |
| *SRC-1* R | CCTCATCCATAAAGCTGTCTCC |
| *ACTR* F | GGGTAAGAGACCGAGGATAGAA |
| *ACTR* R | CAGAGTGGTGGGTATGGAATG |
| *TAFII250* F | GTTCTGAGCTGCCAAGAGATA G |
| *TAFII250* R | CTTCCTCCAGGTTGAGCATAAA |
| *TFIIIC90* F | CATCCTCCTTCCTTGCGTTAT |
| *TFIIIC90* R | CCTGTCCTCATACAGCTTTCC |

**Table S2**: Bacterial strains used in this study

| **Genus and Species** | **Strain** | **Description** | **Source** |
|---|---|---|---|
| *Campylobacter jejuni* | DRH212 | Wild-type *C. jejuni* | This study |

| | | | |
|---|---|---|---|
| *Campylobacter jejuni* | DRH212-pECO102 | *C. jejuni* harboring empty plasmid pECO102 | This study |
| *Campylobacter jejuni* | DRH212 Δ*sliP* | *sliP* deletion strain of *C. jejuni* | This study |
| *Campylobacter jejuni* | DRH212 Δ*sliP*-p*sliP* | *sliP* deletion strain of *C. jejuni* complemented with pECO102-*sliP* | This study |
| *Campylobacter jejuni* | DRH212 Δ*sliP*-p*sliP*$_{G26A}$ | *sliP* deletion strain of *C. jejuni* complemented with pECO102-*sliP*$_{G26A}$ | This study |
| *Campylobacter jejuni* | DRH212 Δ*flgE* | *flgE* deletion strain of *C. jejuni* | (Barrero-Tobon and Hendrixson, 2014) |
| *Escherichia coli* | C3013-pQE30 | *Empty vector expression E. coli strain* | This study |
| *Escherichia coli* | C3013-pQE30-*sliP* | His tagged *sliP expression E. coli strain* | This study |
| *Escherichia coli* | C3013-pQE30-*sliP*$_{G26A}$ | His tagged *sliP*$_{G26A}$ *expression E. coli strain* | This study |
| *Escherichia coli* | DH5α-pCYA-ankB | *E. coli* strain expressing a *Legionella pneumophila ankB* gene fused with an N-terminal adenylate cyclase *cyaA* | (Price et al., 2021) |

| | | | |
|---|---|---|---|
| *Campylobacter jejuni* | DRH212-p*sliP*-*cyaA* | *C. jejuni* strain harboring pECO102 encoding *sliP*-*cyaA* fusion | This study |
| *Campylobacter jejuni* | DRH212-p*ciaB*-*cyaA* | *C. jejuni* strain harboring pECO102 encoding *ciaB*-*cyaA* fusion | This study |
| *Campylobacter jejuni* | DRH212-p*heuR*-*cyaA* | *C. jejuni* strain harboring pECO102 encoding *sliP*-*cyaA* fusion | This study |
| *Campylobacter jejuni* | DRH212-*cobB*::hawkeye | *C. jejuni* strain harboring a transposon insertion in *cobB* | This study |

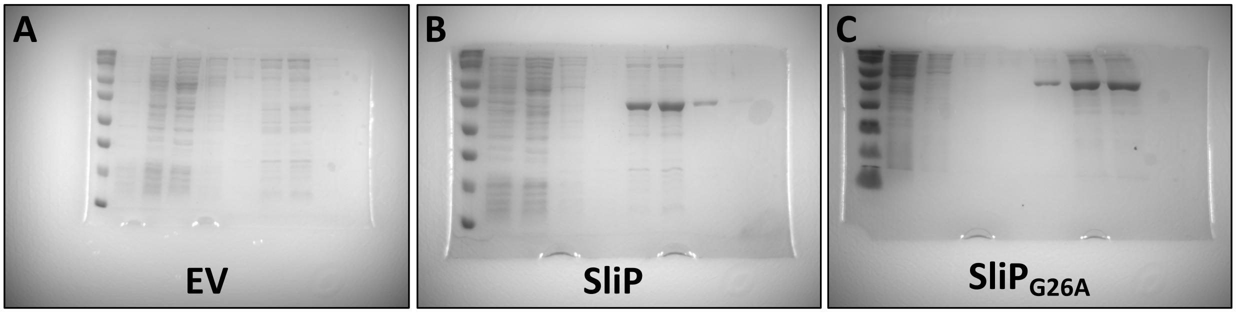


**Figure S1**. Gels of the elutions for the (A) pQE-30 empty vector (EV), (B) SliP, and (C) $SliP_{G26A}$. After packing the Ni-NTA column with lysed *E. coli* supernatant, proteins were eluted from the column. SliP and $SliP_{G26A}$ can be visualized as the predicted molecular weight is 37 kDa.

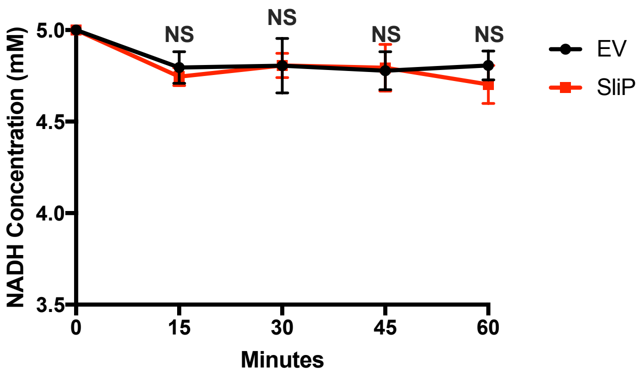


**Figure S2**. Inability of SliP to hydrolyze NADH. To determine if SliP could hydrolyze NADH during deacetylation, 5mM NADH was added to lysine deacetylation assays as previously described. Every 15 minutes, reactions were measured for the abundance of NADH. Multiple comparison testing was performed using ANOVA with a *post-hoc* test.

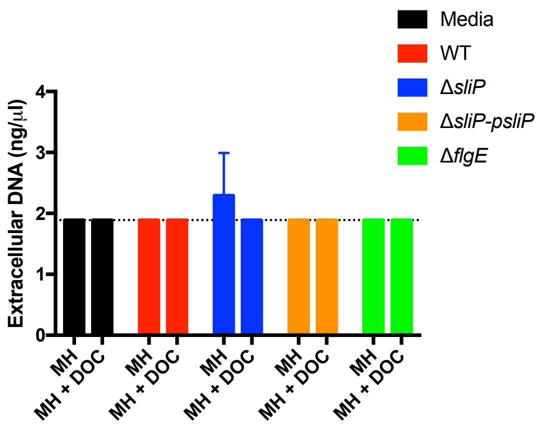


**Figure S3**. Abundance of *C. jejuni* gDNA within MH and MH supplemented with deoxycholate (DOC). After 48 hours of growth in the respective media, the supernatant was isolated and the DNA was extracted as a proxy for cell lysis. Abundance of gDNA was determined using primers specific for *mapA* through qPCR, with most concentrations solely detected at the limit of detection.

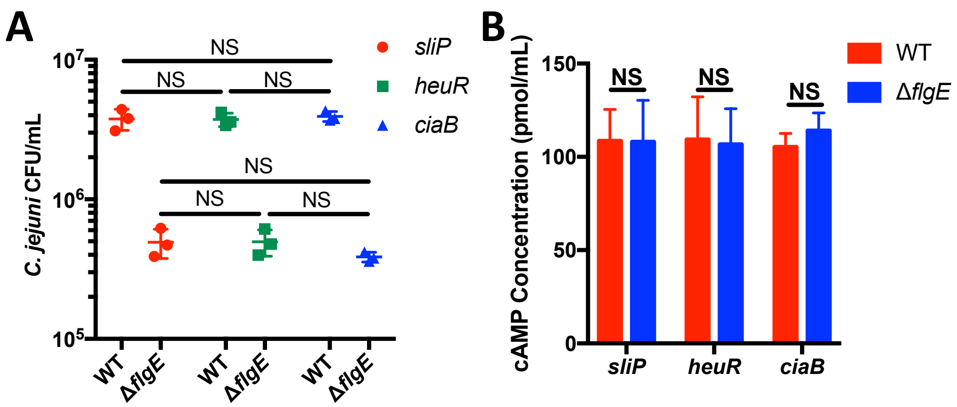


**Figure S4**. CyaA reporter assays do not influence *C. jejuni* internalization within neutrophils. (A) Gentamicin protection assay of *C. jejuni* wild-type and Δ*flgE* harboring the various cyaA reporters for *sliP*, *heuR*, and *ciaB*. (B) *In vitro* CyaA cAMP assay determined that the various strains of reports produce similar levels of functional CyaA. Cell lysates of the various reporter strains were produced and incubated with ATP and recombinant human calmodulin for one hour at 37°C. After incubation, cAMP levels were measured as previously described. Levels of cAMP were not statistically significantly different across the reporters, indicating all strains produce similar levels of functional intracellular CyaA. Multiple comparison testing was performed using ANOVA with a *post-hoc* test.

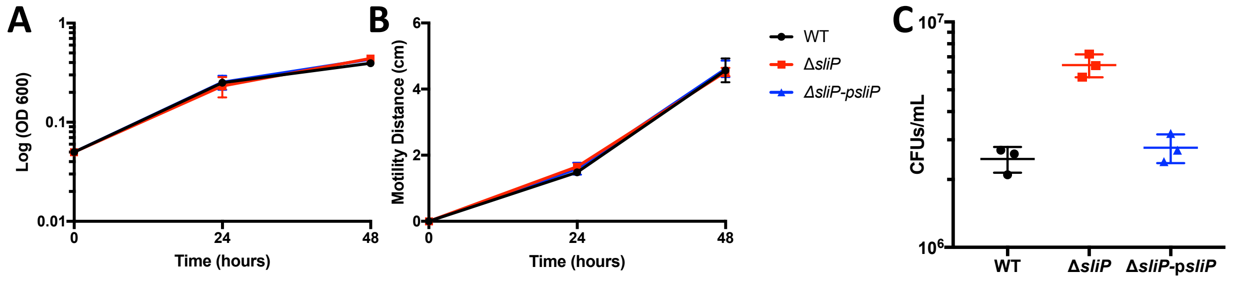


**Figure S5**. Key virulence factors of *C. jejuni* are maintained in the Δ*sliP* mutant. (A) Growth of both strains in MH broth are not significantly different for up to 48 hours. (B) Flagellar motility of both strains in low agar media are not significantly different after 48 hours post inoculation. After 48 hours, the diameter of motility was measured. (C) Intracellular levels of *C. jejuni* within neutrophils are maintained after three hours post-infection. After three hours of incubation, a gentamicin protection assay was performed to determine the abundance of intracellular bacteria. Multiple comparison testing was performed using ANOVA with a *post-hoc* test. ***$p < .001$

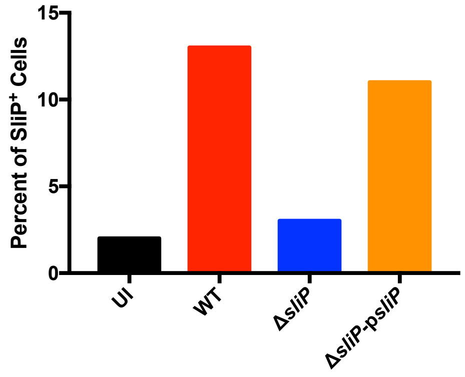


**Figure S6**. Incidence of SliP within uninfected, WT, Δ*sliP*, and sliP complement infected neutrophils. Intracellular abundance was determined using 100 randomly selected cells from each treatment.

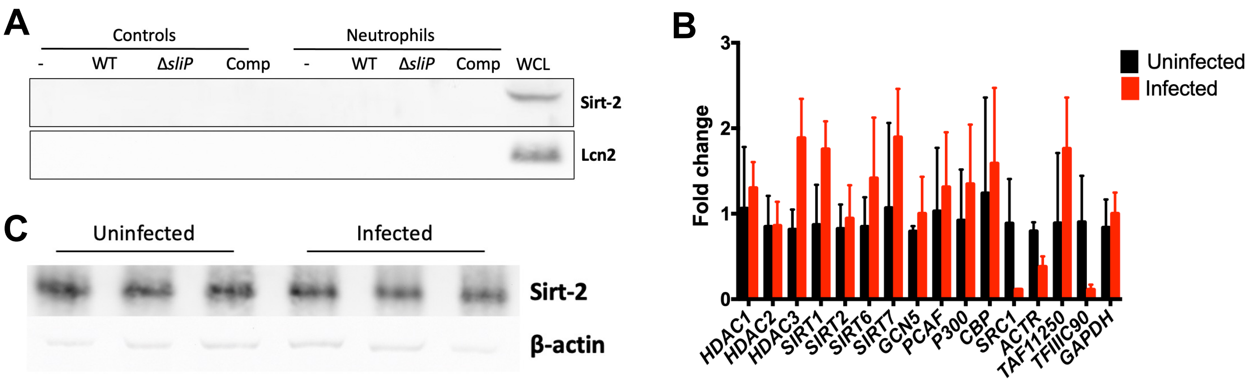


**Figure S7**. Abundance of host sirtuins and histone deacetylases are not upregulated during *C. jejuni* neutrophil infections. (A) Host sirtuin-2 does not translocate and bind to histone H3 during *C. jejuni* infection of neutrophils. Samples from the neutrophil co-immunoprecipitation were probed for the abundance of Sirt-2 as this is a common feature during other bacterial infections. Samples were further analyzed for non-specific cytoplasmic contamination through probing for lipocalin-2 (Lcn2). Whole cell lysates for each protein were included in the blots to ensure the immunoblotting was performed correctly. (B) To determine if histone deacetylation was due to upregulation of a host protein that deacetylates host histones, transcript abundance was determined for proteins that classically target histone H3. Transcript abundance for each gene was normalized to GAPDH abundance. (C) Protein abundance of host sirtuin-2 is not significantly different between uninfected and *C. jejuni* infected neutrophils. Densitometry of Sirt-2 was normalized to the abundance of β-actin.

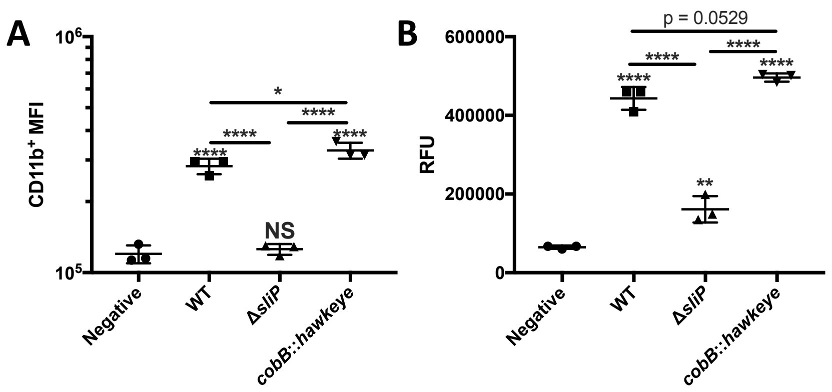


**Figure S8**. The intracellular sirtuin, *cobB*, in *C. jejuni* is not involved in neutrophil activation or NETosis. (A) Neutrophil activation through CD11b expression was significantly increased in the transposon mutant of *cobB* (*cobB*::*hawkeye*) compared to wild-type infected neutrophils. (B) NET abundance through SYTOX staining in the *cobB* mutant was not significantly different from wild-type infected neutrophils. Multiple comparison testing was performed using ANOVA with a *post-hoc* test. *$p < .05$; **$p < .01$; ***$p < .001$; ****$p < .0001$

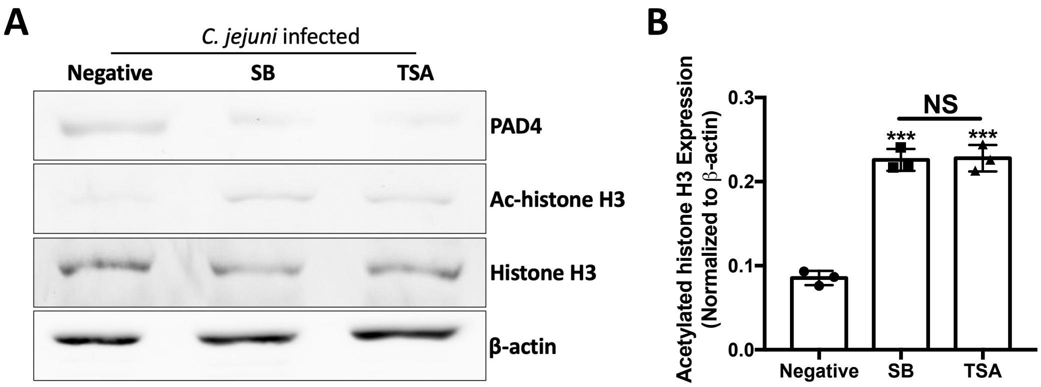


**Figure S9**. Sodium butyrate (SB) and trichostatin-A (TSA) reverse SliP-dependent changes in PAD4 and histone acetylation. Prior to *C. jejuni* infection, neutrophils were pre-incubated with 10mM SB, 100 µM TSA, or media alone (negative) as these concentrations chemically inhibit SliP-dependent lysine deacetylation. SB and SB both significantly reduce the abundance of PAD4 in neutrophils, while additionally increasing the abundance of histone H3 acetylation. PAD4 and histone H3 acetylation abundance was normalized to β-actin and histone H3, respectively.

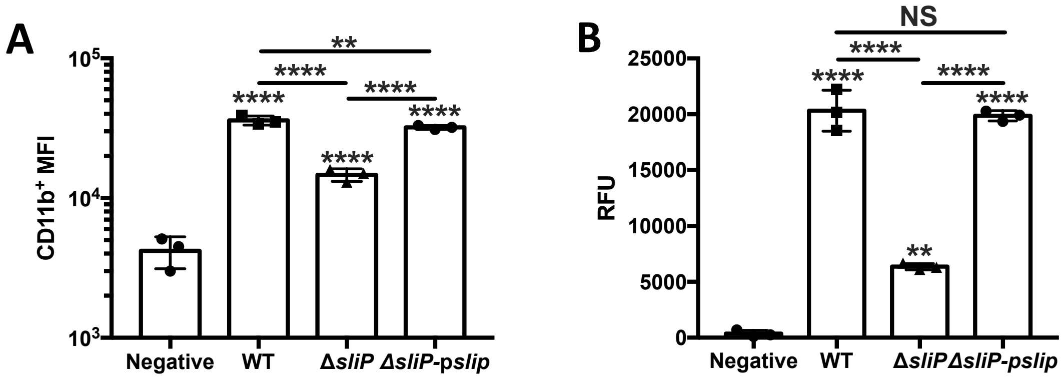


**Figure S10**. *In vitro* murine neutrophil activation and NETosis are SliP-dependent. (A) Neutrophils from C57BL/6 mice were potently activated by wild-type *C. jejuni*, with a significant reduction in CD11b expression compared to wild-type infected neutrophils. Neutrophil activation was restored with the complementation of plasmid-borne *sliP*. (B) NET induction through SYTOX staining was robustly produced, with a reduction in the amount produced by Δ*sliP* infected neutrophils. Complementation through plasmid-borne sliP restored the NET phenotype. Multiple comparison testing was performed using ANOVA with a *post-hoc* test. ***p* < .01; ****p* < .001; *****p* < .0001

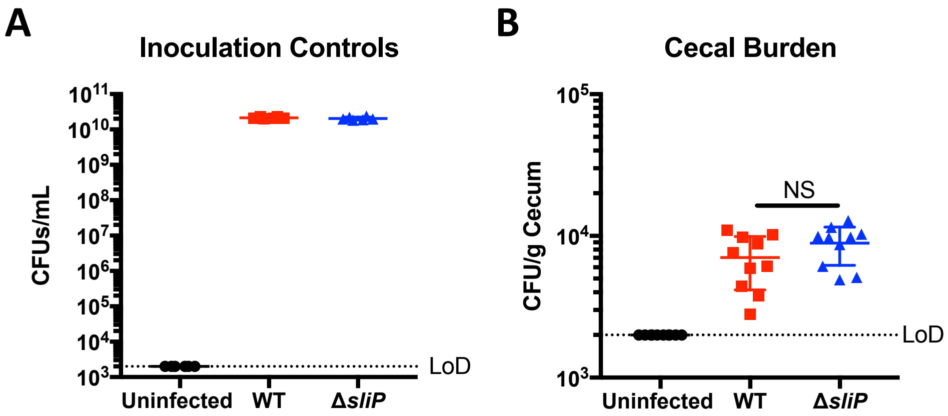


**Figure S11.** Colonization of *C. jejuni* and the Δ*sliP* mutant within the ceca of infected IL-10-/- C57BL/6 mice ten days post-infection. (A) Inoculation control for two separate experiments to ensure mice were inoculated with the correct amount of *C. jejuni*. (B) Cecal burden of *C. jejuni* within uninfected, wild-type, and Δ*sliP* infected mice. Multiple comparison testing was performed using ANOVA with a *post-hoc* test.

**Supplemental Materials and Methods**

Cell lysis dependent genomic DNA quantification

To determine if SliP presence in the supernatant of *C. jejuni* grown in the presence of deoxycholate, concentration of genomic DNA was determined as a proxy for cell lysis. After 48 hours of incubation in Mueller–Hinton (MH) with or without deoxycholate (DOC), supernatant was pulled off for DNA extraction. Before extraction, human DNA was spiked in to account for loss of DNA during the extraction process. For the DNA extraction, one-tenth volume of ice-cold 3M sodium acetate was added to each tube on ice. Next, three volumes of 95% ethanol were added to the tubes on ice. The tubes were then incubated at -20°C for 30 minutes. After incubation, the tubes were centrifuged at 15,000 rpm for 15 minutes at 4°C. After centrifugation, the supernatant was aspirated off, whereby the pellets were washed with 70% ethanol. The tubes were then centrifuged at 15,000 rpm for 5 minutes at 4°C. The supernatant was aspirated off and the DNA pellet were air-dried before resuspension in nuclease-free water. C. jejuni genomic DNA was amplified using mapA-specific primers, while human DNA was amplified using GAPDH-specific primers. Concentrations of each were determined using known DNA concentrations of each alongside the qPCR amplifications.

Gentamicin protection assay

A gentamicin protection assay was performed as previously described (Callahan et al., 2020). Briefly, $10^6$ neutrophils were infected with *C. jejuni* at an MOI of 10 and incubated under microaerobic conditions at 37°C for one hour. After incubation, cells were treated with 100 µl RPMI 1640 containing 10% FBS and gentamicin sulfate for one hour at 37°C under microaerobic conditions. After gentamicin treatment, neutrophils were washed three times with 1x PBSbefore being lysed with 0.1% Triton X-100 for 5 minutes. Cell pellets were resuspended in 1x PBS and serially diluted before being plated on *Campylobacter*-selective media. Cultures were grown for 2 days at 37°C under microaerobic conditions and CFUs were counted. Statistical analysis was performed using unpaired *t* tests and significance inferred at $p < .05$.

*C. jejuni* growth curve

To determine if the *sliP* mutant has a growth defect, bacteria were grown up on *Campylobacter*-specific media for 48 hours under microaerobic conditions. After 48 hours, cells were resuspended in MH broth and normalized to an optical density (OD) of 0.05. After normalization, cells were plated in 96 well plates and incubated under microaerobic conditions at 37°C. Bacterial growth was checked at 24 and 48 hours post-inoculation to determine the growth kinetics. OD readings were performed through a BioTek Synergy microplate reader.

*C. jejuni* motility assay

To determine if the *sliP* mutant has a motility defect, bacteria were grown up on *Campylobacter*-specific media for 48 hours under microaerobic conditions. After 48 hours, cells were resuspended in MH broth and then normalized to an OD of 1 and a final volume of 1 mL. After normalization of all strains, the blunt ends of inoculation loops were dipped into the bacterial culture before being stabbed into low agar MH media. After puncturing the bacterial culture into the media, the inoculation site was indicated. At 24 and 48 hours post-inoculation, the diameter of the zone of motility was determined.

Purification of mouse leukocytes

To obtain murine leukocytes, blood was drawn by cardiac puncture. Neutrophils were isolated by density gradient centrifugation and ammonium-chloride-potassium (ACK) osmotic red blood cell lysis (Akhtar et al., 2010). Briefly, clotted blood was added to 1 mL 1x PBS and centrifuged at 400 x G for 5 minutes. After centrifugation, pellets were resuspended in 500 µl ACK buffer and incubated for 30 seconds before 500 µl 1X PBS was added. Cells were centrifuged at 400 x G for 5 minutes. Cells underwent further ACK lysis treatment until solely leukocytes were obtained. Once isolated, leukocytes were resuspended in RPMI 1640 containing 10% FBS.
